# Pre-amyloid cognitive intervention preserves brain function in aged TgF344-AD rats, maintaining connectivity and enhancing plasticity in a sex-specific manner

**DOI:** 10.1101/2025.03.26.645539

**Authors:** Julia Casanova-Pagola, Federico Varriano, Xavier López-Gil, Genís Campoy-Campos, Elisa López-Bravo, Clara García-González, Enric Abellí-Deulofeu, Raúl Tudela, Emma Muñoz-Moreno, Fernando Aguado, Alberto Prats-Galino, Laura Molina-Porcel, Cristina Malagelada, Guadalupe Soria

**Affiliations:** Brain Connectivity and Neuroimaging Lab, Institute of Neurosciences, Faculty of Medicine and Health Sciences, University of Barcelona, Carrer de Casanova 143, 08036 Barcelona, Spain; IDIBAPS, Magnetic Resonance Imaging Core Facility, Carrer del Rosselló 149-153, 08036 Barcelona, Spain; Biomedicine Department, Institute of Neurosciences, Faculty of Medicine and Health Sciences, University of Barcelona, Carrer de Casanova 143, 08036 Barcelona, Spain; CIBER de Bioingeniería, Biomateriales y Nanomedicina, Instituto de Salud Carlos III, Avenida Monforte de Lemos 3-5, 28029 Madrid, Spain; Department of Cell Biology, Physiology and Immunology, Biology Faculty, University of Barcelona, Carrer Diagonal 643, 08028 Barcelona, Spain; Laboratory of Surgical Neuroanatomy (LSNA), Faculty of Medicine and Health Sciences, Universitat de Barcelona, Barcelona, Spain; Alzheimer’s disease and other cognitive disorders unit. Neurology Service, Hospital Clínic, I Fundació de Recerca Clínic Barcelona-Institut d’Investigacions Biomèdiques August Pi i Sunyer (FRCB-IDIBAPS) and University of Barcelona, Carrer del Rosselló 149-153, 08036 Barcelona, Spain; Neurological Tissue Bank, Biobanc-Hospital Clínic-FRCB-IDIBAPS, Carrer del Rosselló 149-153, 08036 Barcelona, Spain; Centro de Investigación Biomédica en Red sobre Enfermedades Neurodegenerativas (CIBERNED), Instituto de Salud Carlos III, Avenida Monforte de Lemos 3-5, 28029 Madrid, Spain

**Keywords:** Alzheimer’s disease, Cognitive Reserve, Cognitive Stimulation, Functional Connectomics, Magnetic Resonance Imaging, Neuroinflammation, Neuronal plasticity, TgF344-AD model, Sexual dimorphism

## Abstract

**Background:** Alzheimer’s disease is characterized by progressive cognitive decline and neurodegeneration, with cognitive reserve playing a key role in mitigating disease impact. Cognitive stimulation has been suggested as a non-pharmacological approach to enhance cognitive reserve and delay cognitive deterioration, but its underlying mechanism remains to be fully understood. This study investigates the effects of pre-amyloid cognitive intervention on brain connectivity, memory, synaptic plasticity and neuroinflammation in aged TgF344-AD rats, considering sex-specific differences.

**Methods:** Male and female TgF344-AD and wild-type rats were assigned to trained and untrained groups, with cognitive stimulation administered through repetitive delayed nonmatch-to-sample tasks. Longitudinal magnetic resonance imaging acquisitions assessed training-induced changes in whole-brain functional connectomics and in particular entorhinal cortex connectivity. Memory was evaluated using the novel object recognition test. Cellular analysis of neurons (NeuN^+^, Parvalbumin^+^) and microglial cells, as well as molecular (PSD95, TrkB, p-RPS6, and VGLUT) analyses were conducted to determine the role of cognitive stimulation in modulating neuronal density, neuroinflammation and neuroplasticity.

**Results:** Male TgF344-AD rats undergoing prolonged cognitive stimulation had preserved global functional connectivity and exhibited improved recognition memory, compared to untrained animals, while TgF344-AD females did not follow this pattern. Entorhinal cortex connectivity was significantly loss in 19-month-old rats compared to wild-type rats and this was completely prevented by training. At a cellular level, cognitive stimulation significantly decreased the number of PV^+^ neurons in the dentate gyrus of trained rats. Moreover, a greater microglial density around Aβ plaques and a less reactive phenotype was clearly observed at 11 in trained rats. These protective effects diminished by 19 months, coinciding with increased neuroinflammation and microglial dysfunction. At a molecular level, cognitive stimulation preserved PSD95 expression in male TgF344-AD and p-RPS6 in both sexes.

**Conclusions:** Pre-amyloid cognitive stimulation enhances synaptic plasticity, sustains brain network integrity, and modulates neuroinflammation, contributing to increased resilience against Alzheimer’s disease-related cognitive decline. In general, cognitive stimulation exerted a more protective effect in male TgF344-AD rats showing sex-dependent differences in pathology and cognitive reserve mechanisms. These findings highlight the importance of early cognitive engagement as a potential strategy to delay disease onset and underscore sex-specific differences in cognitive resilience mechanisms.

## 1. Background

Alzheimer’s disease (AD) is a neurodegenerative disorder that accounts for the main cause of dementia worldwide. It is characterised by a progressive and irreversible decline in cognitive function and impaired hippocampal-dependent learning and memory. Currently, over 50 million people suffer from dementia globally and its prevalence is predicted to triple by 2050 (1). In the absence of effective treatments, early detection, advances in diagnosis and the applications of treatments that can delay symptomatic presentation are urgent.

Substantial evidence shows that socio-behavioural proxies such as years of education, activities of daily living, physical exercise and environmental enrichment are associated with less cognitive decline, cognitive flexibility and reduced risk of dementia in AD, possibly by increasing the threshold at which deterioration manifests clinically (2–4). These engaging activities contribute to brain health and cognitive reserve (CR), which is the ability to maintain well-preserved cognitive function despite the presence of brain deterioration, promoting resilience to pathology (2,5). The concept of CR postulates that enriching experiences are functionally protective against damage, as the brain actively copes with or compensates for pathology by either using pre-existing cognitive processing or utilising compensatory mechanisms (6,7). This protective effect can arise from interacting factors such as increased neuronal plasticity, more efficient synaptic connections and modulating neuroinflammatory process, all culminating in enhanced CR and capacity to cope with pathology and preserve neuronal function (8,9). Therefore, it is crucial to investigate mechanisms to increase CR, since it is a promising therapeutic approach to delay dementia onset. Indeed, delaying the onset of dementia by 1 year is estimated to reduce age-dependent dementia by more than 10% (10).

Given the late onset of symptomatology and the difficulty to identify subjects in the early stages of the disease, research using transgenic animal models of AD provides a useful tool to improve our understanding of the mechanisms and identify potential novel targets to treat AD. In this line, the highly translational TgF344-AD (TG) rat model of AD recapitulates the main hallmarks of AD pathology in an age-dependent manner, as observed in patients. Specifically, TG rats develop progressive cerebral Aβ accumulation, preceding tauopathy, gliosis, neuronal death and cognitive impairment (11).

In the past, we have used non-invasive magnetic resonance imaging (MRI) techniques to characterize the changes in functional and structural global brain connectivity in this TG rat model. Alterations in structural and functional brain network properties were described at early ages (5–6 months) (12). Moreover, aging significantly affected structural network metrics in TG rats, while whole-brain network metrics describing integration and segregation of the functional connectomes remained unaffected (13). We hypothesized that training and repetition of the delayed non-match to sample (DNMS) task, used to evaluate cognitive skills, could have induced a learning effect, increasing the CR of these animals and therefore preserving their functional connectivity. Indeed, we showed that structural brain networks impacted the cognitive outcome of TG rats, while functional connectivity correlated with cognitive performance only in the WT group. Moreover, resting state networks were investigated in the same cohort and significant alterations in somatosensory and default mode networks were found only at early (5 months) or late stages (18 months), supporting the idea that the repetition of the DNMS task, contributed to maintain cognitive performance, at least until 15 months of age (14). This type of cognitive intervention may increase CR by modifying the evolution of functional networks and their connections in TG rats as a compensatory mechanism, at least during the intermediate stages.

The DNMS task has conventionally been used to evaluate working memory, which relies on the hippocampus, as well as the prefrontal and entorhinal cortices as major components of memory-related functions (15,16). It requires flexibility in behaviour throughout time as it is a complex, repetitive task that engages with working memory, a functionally important system underlying higher order domains, which has been shown to correlate significantly with IQ in humans (15,17). Based on our previous observations and the inherent challenge that the DNMS task presents, in our study we used it as a form of cognitive stimulation. In the same line, previous studies have used other hippocampal-based repetitive tasks, such as the Morris water maze as a form of cognitive training in AD murine models (18,19).

The study of CR requires a comprehensive approach that integrates measures of brain pathology, cognitive performance, and factors influencing this relationship. According to recent guidelines, CR studies should include (1) an assessment of brain changes due to aging or disease, (2) measures of cognitive function, and (3) an identified proxy or mechanism that moderates the impact of brain pathology on cognition (7). In our study, we apply these principles to investigate whether intensive training and repetitive testing of DNMS through life increases CR. Thus, we assessed brain connectivity using MRI-based metrics, ensuring that changes in network integrity are objectively measured. Additionally, we evaluated memory performance through the novel object recognition (NOR) test, allowing us to establish a link between brain function and cognitive outcomes. At the cellular level, we examined number of neurons and parvalbumin positive interneurons in the hippocampus. As synaptic integrity is a key marker of CR mechanism, synaptic-related proteins were analysed at a molecular level. Furthermore, we investigated the role of microglia, measuring both their morphology and density around Aβ plaques, as neuroinflammatory responses are increasingly recognized as mediators of CR in neurodegenerative conditions. Importantly, we include both female and male subjects to examine sex differences in CR mechanisms, addressing a crucial yet often overlooked factor in preclinical research.

## 2. Methods

### 2.2. Animals and experimental design

Male and female TgF344-AD rats (30) and wild-type (WT) Fischer rats (30) were housed under controlled conditions of temperature (22 ± 1°C) and humidity (55 ± 10 %) on a 12 h light/12 h dark cycle. Water and food were available *ad libitum* except during behavioural tests. Animal work was performed in accordance with the local legislation (Decret 214/1997 of July 30^th^ by the Departament d’Agricultura, Ramaderia i Pesca de la Generalitat de Catalunya) with the approval of the Experimental Animal Ethical Committee of the University of Barcelona, and in compliance with European legislation.

MRI scans were performed at 5 time points: 3, 7, 11, 15 and 19-months-old, with the first scan being performed before DNMS training. Rats were then divided into two groups: untrained and cognitively trained with the DNMS task. At 3 months of age, rats began the habituation protocol and the DNMS training. At 7 months old, rats performed the DNMS task for 10 consecutive days every 4 months. The novel object recognition (NOR) test was performed at 19 months, after which rats were sacrificed and their brains extracted for histological and protein assays. A small subgroup of rats (n=3-4 per experimental group) was sacrificed at 11 months to study the biological effects of cognitive training in middle adulthood (Fig. 1). A specific breakdown of the number of animals used in each experiment can be found in the supplementary table 1.

**Figure 1.**
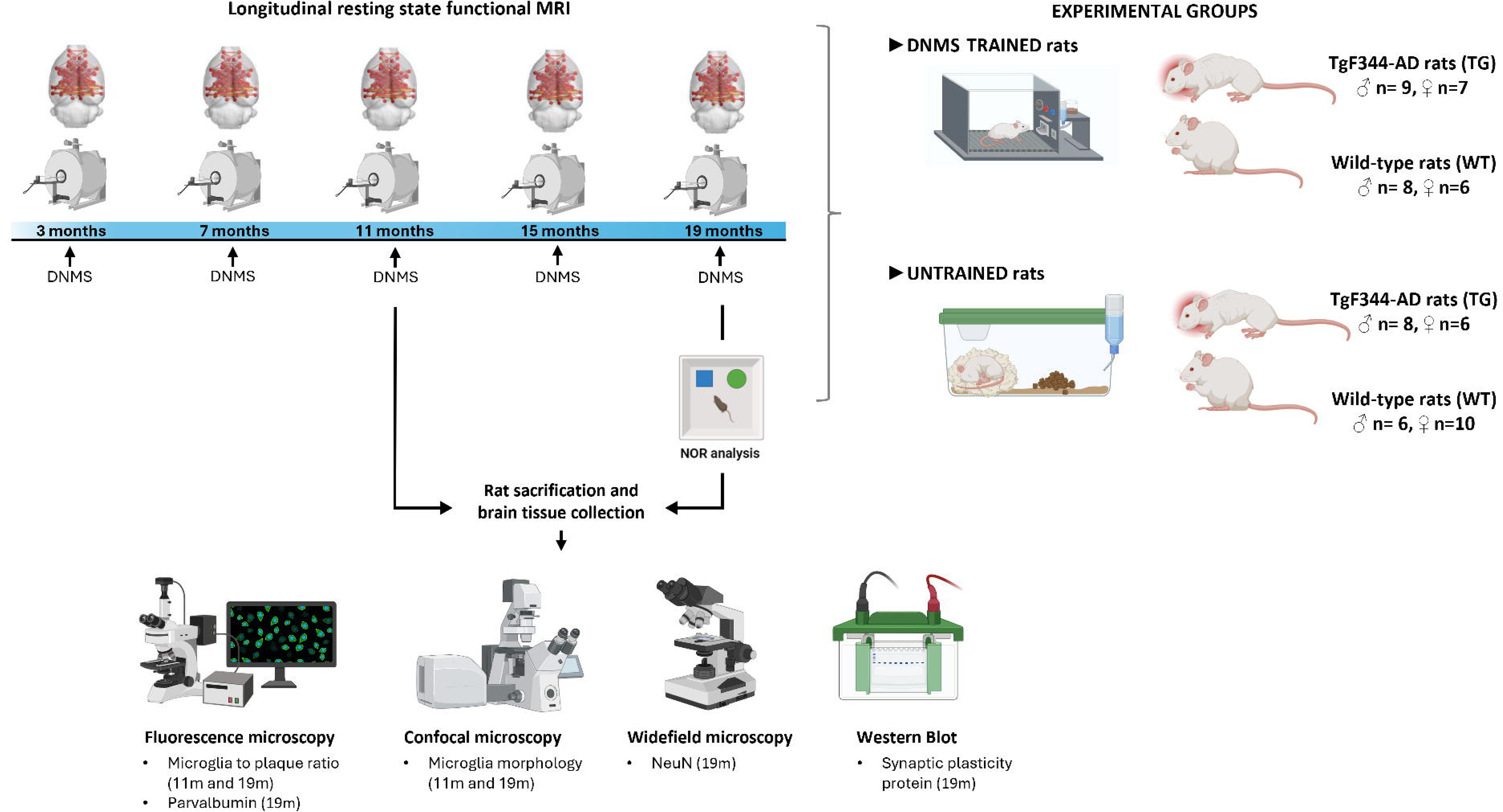
Schematic representation of the experimental timeline. All animals underwent an initial MRI scan at 3 months old. Then, they were divided into four experimental groups, WT and TG with or without training and periodic execution of the DNMS task. Follow-up scans were performed in all groups at 7, 11, 15 and 19 months. The NOR test was performed at 19 months to evaluate the recognition memory. A reduced subset of animals was sacrificed at 11 months (n= 4-5 per experimental groups) while the remaining subjects were sacrificed at 19 months. In both cases, brains were extracted for histological and biochemical analyses. Figure created with BioRender.

### 2.3. Behavioural paradigms

#### 2.3.1. Training and Delayed Non-Match to Sample task

Cognitive stimulation by DNMS task was performed to enhance CR. The DNMS task was performed in isolated operant chambers (Med Associates, Fairfax, VT, USA) containing 3 extendable levers: two placed on the pellet wall (right and left levers) and the third, centre-placed on the opposite wall. Rats underwent habituation, training and task phases as previously described (12). In brief, the habituation phase involved exposing the rats to 5 minutes of daily handling for 7 days and placing them in the DNMS chambers for 30 minutes. The DNMS training phase consisted of six levels of ascending difficulty. Once the acquisition criteria was achieved, the DNMS task began (12). During testing weeks, the animals received 75% of their standard food intake, to increase their performance motivation. The DNMS task started with the sample phase, where the animals had to press the lever that extended (left or right). After a random 1 to 30-second wait, the centre-placed lever on the opposite wall would extend, and the rats had to press it. If successful, both right and left levers would extend again, and the rats had to press the lever which had not been presented during the sample phase (non-matched) to obtain a sucrose pellet reward. An incorrect response (pressing the matched lever) led to a 5-second time-out with overhead lights turned off and no pellet reward. The DNMS tasks were finalized after 60 minutes or when 90 trials were performed.

#### 2.3.2. Novel Object Recognition Test

The NOR test was used to evaluate the rat’s working memory. At 19-months-old, rats were individually placed in empty testing arenas (40 cm x 40 cm x 30 cm) for 5 minutes for habituation purposes. The familiarisation and testing phase were performed the following day, where rats were placed in the testing arena for 5 minutes with two identical objects. After a 30-minute delay, rats were placed back in the arena with the following two objects: a familiar one (from the familiarisation phase) and a novel object. Interactions with these objects were recorded for 5 minutes with a digital video camera mounted overhead. The SMART 3.0.06 video tracking software was used to analyse the recordings and track the time each rat spent exploring each object, to quantify the recognition index (RI), the time spent interacting with the novel object relative to the time spent interacting with both, as well as the total exploration time. Object exploration was counted when rats were within a 2 cm perimeter of an object, with their snout directed towards it. For analysis only the first 2.5 minutes of each 5-minute recording were used, as the rats lost interest and showed reduced locomotor activity over time, potentially masking cognitive deficits between experimental groups (20).

### 2.4. Magnetic Resonance Imaging

MRI experiments for functional connectomics were performed as previously described on a 7.0-T BioSpec 70/30 horizontal animal scanner (Bruker BioSpin, Ettlingen, Germany) equipped with an actively shielded gradient system (400 mT/m, 12-cm inner diameter) (15). Briefly, animals were placed in a prone position in a Plexiglas holder with a nose cone for administering anaesthetic gases (1.5% isoflurane in a mixture of 30% O2 and 70% CO) and were fixed using tooth and ear bars and adhesive tape. To ensure stability during the resting state functional magnetic resonance imaging (rsfMRI), rats received a 0.5 ml bolus of medetomidine (0.05 mg/kg; subcutaneously (s.c.)) and a catheter was implanted in their back for continuous perfusion of medetomidine. Isoflurane was gradually decreased to 0.5% and 15 minutes after the bolus the medetomidine perfusion (0.1 mg/kg/h; s.c.) started at a rate of 1ml/hour. The acquisition protocol included the following imaging: T2-weighted images, acquired using a rapid acquisition with relaxation enhancement (RARE) sequence with effective echo time (TE) of 35.3 ms, repetition time (TR) of 6000 ms, RARE factor = 8, voxel size = 0.12x0.12 mm², 40 slices, slice thickness = 0.8 mm and field of view (FoV) = 30x30x32 mm³; T1-weighted images, acquired using an Modified Driven Equilibrium Fourier Transform (MDEFT) protocol with TE = 2 ms, TR = 4000 ms, voxel size = 0.14x0.14x0.5 mm³ and FoV = 35x35x18 mm³. For rsfMRI, a gradient echo T2* acquisition was used, with the following parameters: TE = 28 ms, TR = 2000 ms, 600 volumes (20 minutes), voxel size = 0.4x0.4x0.6 mm³, FoV = 25.6 x 25.6 x 20.4 mm³.

#### 2.3.1. Image processing and connectome definition

The acquired images were processed to obtain the functional connectomes. Briefly, at each age timepoint, a T-2 weighted group template was constructed by interactive multi-stage registration using ANTs (21), and a modified SIGMA rat brain atlas was registered to the corresponding group template (22). Then the atlas parcellation and segmentation were registered to each subject’s T2-weighted images using the group template as an intermediate registration step to obtain individual brain masks and region parcellations. RsfMRI volumes were denoised, skull-stripped and intensity-normalised using MRTrix3 (23). Atlas parcellation and segmentation were registered from each subject’s T2 space to their respective pre-processed rsfMRI spaces to define the regions between which connectivity was assessed.

RsfMRI processing was done with Nilearn (24). Time series were detrended and standardised, a band-pass filter was applied to keep frequencies between 0.01-0.1Hz and a gaussian smoothing kernel was applied with a full width at half maximum of 1mm to enhance signal-to-noise ratio and increase statistical sensitivity.

Since brain activity identified by rsfMRI has been constrained to GM [(25) only regions comprising grey matter tissue (94 regions) were considered as nodes in the functional connectome. The connection weight between nodes was calculated as the partial correlation between the pair of regional time series, converted to z-scores by applying Fisher’s z-transformation. Since correlations are symmetric, undirected graphs will be constructed. Negative correlation coefficients were excluded since negative functional connectivity have potentially artefactual origins and unclear physiological interpretation. All the connections with positive weight (z>0) were considered for weighted undirected functional connectome construction. Binary undirected functional connectomes were constructed by setting to 1 all non-zero values of their weighted counterparts.

#### 2.4.1. Connectome metrics

Brain network organisation was described using graph theory, taking different brain regions as nodes and functional connectivity as vertices. This allows us to study functional connectivity at the network level by capturing the complex and dynamic interplay between brain regions in a systematic and quantifiable manner. Specially we focused on global efficiency and clustering coefficient. These metrics provide a description of integration and segregation at a global level (26)[. Global efficiency measures network integration: the ability to combine information from different regions. It is inversely related to the shortest path length and it is calculated as the average efficiency between all pairs of nodes. Higher global efficiency characterises stronger and faster communication through the network. Network segregation, the ability for specialized processing within densely interconnected groups of regions, was quantified by the clustering coefficient. The average clustering coefficient is the average of nodal clustering coefficients, which are calculated as the fraction of possible triangles that exist between each node and its neighbours (triangles being sets of three nodes that are all mutually connected by edges). High values of clustering coefficient are related to highly segregated and connected networks. These metrics have been commonly used in human studies of AD (27–30), and therefore, can provide comparable and translational results. For longitudinal functional network metrics comparison between groups, a total of 5 data points from different subjects were discarded due to MRI acquisition and processing issues.

#### 2.4.2. Seed-based analysis of the Entorhinal Cortex

Seed-based analysis is a model-based method in which a region of interest (ROI) is selected as a seed and correlations between its average blood oxygen level-dependent (BOLD) signal and the BOLD signal of all brain voxels are calculated (31). We performed seed-based analyses using the left entorhinal cortex (EC) as seed, segmented from the rat brain MRI atlas (22) to build a seed-based functional connectivity map, providing information on which brain regions are functionally connected. Correlation maps were calculated using the Nilearn package and were converted to z-scores using Fischer’s transformation and a non-parametric permutation inference procedure (32) was applied to determine which brain regions showed different connectivity patterns between groups by implementing a non-parametric t-test with a total of 5000 permutations per contrast. The resulting p-values maps were then processed to detect cluster-like structures using the Threshold-Free Cluster Enhancer (TFCE) procedure (33) and multiple comparisons were adjusted by controlling the Family-Wise Error (FWE) Rate ((α=0.05). Images were visualised using the ITK-SNAP software (version 4.2.0) (34).

### 2.2. Tissue preparation

At 11 (n=16) and 19 (n=43) months, after the corresponding MRI session with rats under deep anaesthesia, transcardiac perfusions were performed with phosphate-buffered saline (PBS) washes and with 4% paraformaldehyde. Rats were decapitated and the brains were extracted. Both hemispheres were separated, one for histological studies and the other was dissected, quickly frozen, and stored at -80°C for biochemical studies. The hemisphere used for histological studies was coronally cut into 6 slices of 3mm, excluding the cerebellum. The slices were mounted on paraffin blocks (Leica EG 1150H) and subsequently sliced into 5 µm sections using a microtome (Leica RM2255).

### 2.3. Immunohistochemical studies

Deparaffinized 5 µm coronal sections were incubated with PTlink at pH 6 (20 minutes at 95°C) and then with 98% formic acid (5 minutes). A peroxidase blocking solution (Dako) incubation was performed (30 minutes) before the 45-minute incubations with primary rabbit anti-NeuN antibodies (Ab) (1:1000; Abcam, #177487) or primary mouse anti-Aβ Ab (1:100; Dako, Clone #6F/3D). For signal amplification rabbit (Dako, #K8009) or mouse linker (Dako, #K8021) were added (30 minutes), prior to incubations with HRP-linked anti-rabbit (Cell Signalling Technology, #7074) or anti-mouse (ThermoFisher Scientific, #32430) secondary antibodies (30 minutes). 3,3’-Diaminobenzidine (DAB) chromogen was added (10 minutes) and slides were stained with haematoxylin. Washes with Dako buffer were performed in between incubations. Tissues were mounted onto slides, dehydrated and covered with coverslips ready for microscopy use. All steps were performed in a humid chamber, at room temperature (unless stated otherwise), in the dark until DAB was added.

### 2.4. Immunofluorescence studies

Deparaffinised coronal sections were incubated with PTlink (20 minutes at 95°C) followed by an incubation with 98% formic acid (5 minutes). LinkerMouse incubation was followed by a 1 hour 5% bovine serum albumin (BSA) incubation and 45-minute incubation with primary mouse anti-Aβ Ab (1:100; Dako, Clone #6F/3D), rabbit anti-Iba1 (1:400; Wako, #019-19741) and rabbit anti-parvalbumin (1:500; Proteintech, #29312-1-AP). Sections were incubated for 1 hour in the dark with polyclonal goat anti-mouse Alexa Fluor 488 (1:400; Thermo Fisher Scientific, #A31620), or polyclonal donkey anti-rabbit Alexa Fluor 546 secondary antibodies (1:400; Thermo Fisher Scientific, #A10040). Sections were rinsed in 70% ethanol for 5 minutes and stained with saturated (70 % EtOH) Sudan Black solution for 15 minutes.

### 2.5. Image acquisition and analysis

For NeuN^+^ cell count and quantification of microglia to plaque size ratio, the BX-51 Biological Binocular Microscope (Olympus) was used. For unbiassed NeuN^+^ count, the optical fractionator technique was applied using the Stereo Investigator Software (Leica). The 20x objective was used to obtain NeuN^+^ images of the areas of the EC and hippocampus (Dentate Gyrus (DG), CA1, CA3). For quantification, the perimeter of each area was delimited as a ROI, and a randomised grid-counting system was used to estimate the total neuronal population per area. For each area, the estimated neuronal population was then normalized to its respective ROI to obtain a NeuN^+^ count/area. To obtain images of microglia around Aβ plaques, the 40x objective was used to select a pre-determined ROI around each plaque per image, and the microglia within each ROI were considered as plaque associated. Images of parvalbumin-positive (PV^+^) cells were acquired using a widefield AF6000 optical microscope (Leica) at 20x magnification. The LAS X software (Leica) was used to obtain mosaics of the hippocampus (DG, CA1, CA3). PV^+^ counts were then performed manually on Image J. Total PV^+^ counts per region were normalised to their respective ROIs to obtain a ratio of PV^+^ counts/area. The confocal SP5 2 photon microscope (Leica) was used to generate 3D images of Aβ^+^ plaques and Iba1^+^ microglial cells in the EC. The rhinal fissure served as the anatomical reference, and images were acquired in a ventral direction. The 40x oil-immersed objective was used to obtain 4 images per rat. Images were 1,024[×[1,024 pixels, obtained at a 400 Hz speed with z-stacks using the LAS AF software. Images of the hippocampus (DG, CA1, CA3) were obtained in - 3.2 mm Bregma, and of the EC in - 5.2 mm Bregma, based on Bregma coordinates from the Paxinos and Watson’s Atlas of Stereotaxic Surgery (35). ImageJ software (version 2.14.0/1.54f; Java 1.8.0_322 (64-bit)) was used to process and analyse images obtained from microscopy. For the analysis of microglial morphology, the macros “macro.ijm” and “composite_macro.ijm” were used. The plugin “AnalyzeSkeleton 2D/3D” was used to binarise and skeletonise images for the quantification of branch numbers. Eight microglia were analysed per subject and the average per rat was analysed and expressed graphically.

### 2.6. Western Blot analyses

Brain cortical segments including the motor, somatosensory, auditory, insular and EC (Bregma 1 to -3.55) were used for the Western Blots. Supernatant for analysis was obtained by defrosting samples with an adapted lysis buffer (36). Samples were then sonicated, incubated for 2 hours at 90°C and 300 rpm and centrifugated at 4°C and 1000 g for 10 minutes, the total concentration of the supernatant was quantified with the protein assay kit (Bio-Rad, #5000114), using BSA for the calibration curve (Bio-Rad, #5000007) and following manufacturer’s instructions. Aliquots were stored at -80°C until use.

Samples obtained from the rats’ cortex (15 – 20 µg of protein) were separated in NuPAGE™Novex™ 4-12% SDS-acrylamide gels and proteins were transferred to nitrocellulose membranes with the iBlot™ dry blotting system (Thermo Fisher Scientific). Primary antibodies were diluted in Tris-buffered saline containing 0.1% Tween-20 and 5% BSA and incubated overnight at 4°C. Secondary Abs were diluted in TBS-Tween with non-fat, dry 5% milk (Bio-Rad) for 1 hour. Supersignal™ West Pico Plus chemiluminescent substrate was used for protein detection and images were obtained using ChemiDoc™ (Bio-Rad). The following primary Abs were used at a 1:1000 dilution: mouse anti-PSD95 (Thermo Fisher Scientific, #MA1-045), rabbit anti-VGLUT1 (Synaptic Systems, #135303), rabbit anti-p-RPS6 (Ser235/236) (Cell Signaling Technology, (CST) #4858), mouse anti-S6 (CST, #2317), and mouse anti-TrkB (BD Biosciences, #610102).

Photoshop (version 25.4.0) and ImageJ were used for image analysis and the quantification of relative protein expression obtained from Western blots.

### 2.7. Statistical Analysis

Analyses were performed using a mixed-effect model, a two-way ANOVA, or three-way ANOVA, depending on the number of factors analysed, as specified in each figure legend. When appropriate, post-hoc Tukey tests were conducted following two-way ANOVA, and Fisher’s least significant difference (Fisher’s LSD) tests were performed following three-way ANOVA. Data was analysed separately by sex except in histological studies where the 11 months-old groups were too small to divide by sex. Data was processed and analysed blindly. Statistical analyses were performed with the GraphPad Prism 10 software (version 10.2.2) and data were presented as mean ± standard error of the mean (SEM), considering p<0.05 as statistically significant. The complete results from the ANOVAs, including F-values, degrees of freedom and p-values, are available in supplementary table 2.

## 3. Results

### 3.1. Cognitive stimulation preserves global functional connectomics in male TG

To determine whether cognitive stimulation by training and repetitive DNMS testing had an effect on brain connectivity and organisation, TG and WT rats underwent a longitudinal MRI study, with acquisitions every 4 months from 3 to 19 months of age (Fig. 2). Since integration and segregation are two fundamental principles of brain organisation, we computed the global efficiency and clustering of the resting state functional connectome. A three-way ANOVA revealed a significant interaction between age and treatment (p<0.05) and a significant main effect of age (p<0.01, p<0.001) and treatment (p<0.001) on the global efficiency and clustering in male rats, respectively (Fig. 2a and 2c). This is indicative of different temporal patterns of integration and segregation aspects of functional connectivity amongst the different treatment groups. Notably, at both 7 and 11 months of age, trained rats, have significantly higher global efficiency compared to their untrained littermates (WT, p<0.05; TG, p<0.01, at both time points) (Fig. 2a). In the case of female rats, a three-way ANOVA revealed a significant interaction between age and genotype (p<0.05) in the global efficiency and a significant effect of age (p<0.05) in both global efficiency and clustering (Fig. 2b and 2d, respectively). Full statistical details are available in table 2 of the supplementary materials.

**Figure 2.**
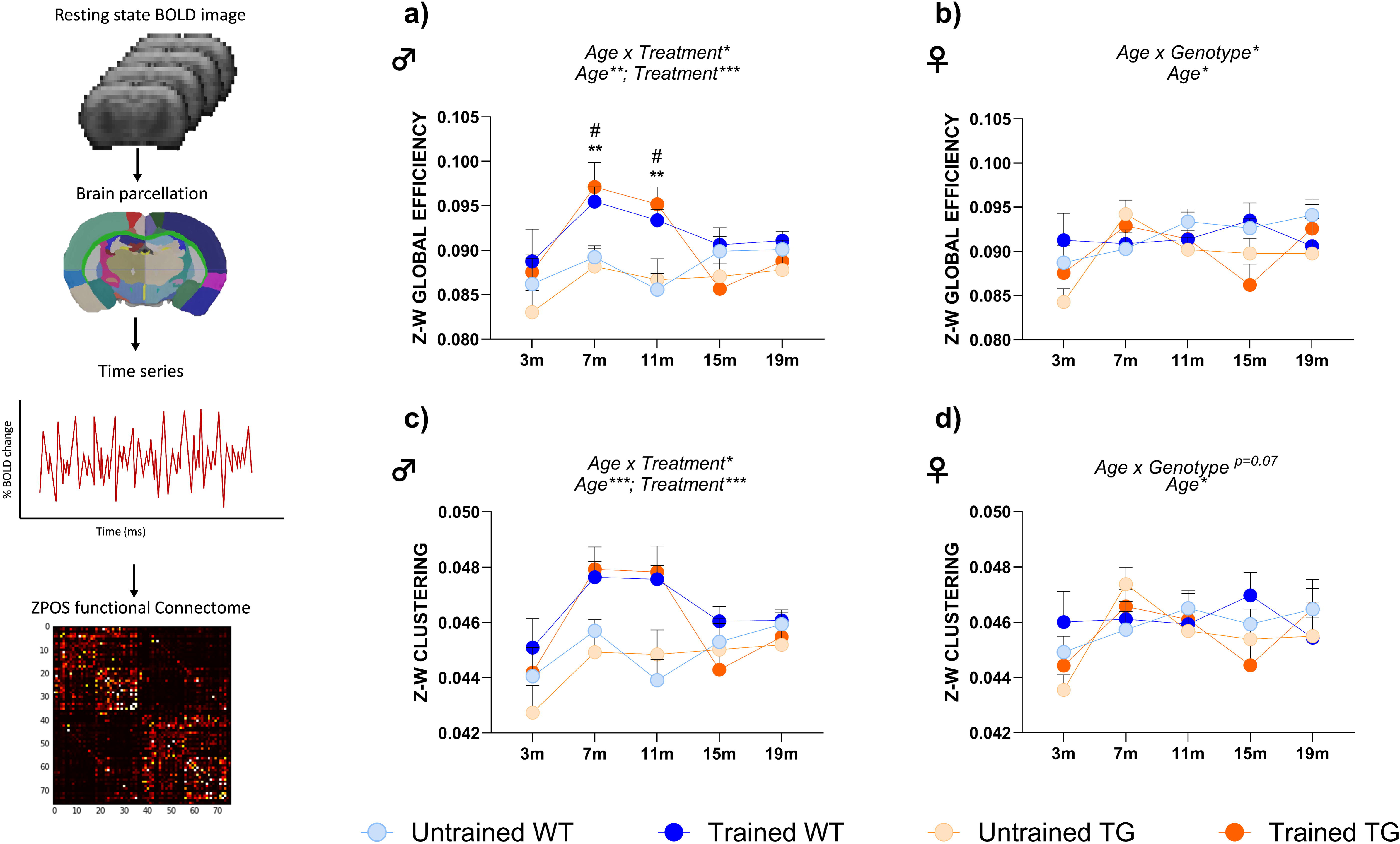
Whole brain functional connectomics are preserved in cognitive stimulated male TG rats. Data shows global efficiency (a and b) and clustering (c and d). The effects of genotype, treatment and age and the interactions between these factors were assessed by mixed-effect analyses and Tukey’s multiple comparisons test. N=5-6 per experimental group. Data is expressed as the average metric value ± SEM * p<0.05; ** p<0.01; *** p<0.001. Significant differences between treatment groups within each time point are denoted with “#” for WT rats and “*” for TG.

### 3.2. Cognitive stimulation restored EC functional connectivity in TG Rats

At 19-months-old, we performed a seed-based analysis of the EC (Fig. 3) due to its function as the gateway of information flow to the hippocampus, its central role in working memory and its early degeneration in AD (37,38). Seed-based analyses demand a larger sample size, which prevented us from disaggregating the data by sex. Therefore, the results are reported with sexes combined. The analysis revealed genotype-specific differences in the functional connectivity maps of untrained rats, as evidenced in the average functional connectivity maps (Fig. 3a and b). These results highlight how brain regions co-activate with the seed differently per genotype. The statistical maps highlight the brain regions where the co-activation was significantly lower in TG rats compared to WT littermates. Regions with altered functional connectivity included the visual, somatosensory, motor, cingulate and retrosplenial cortices (WT > TG; p<0.05, FWE-corrected) (Fig. 3a). These baseline connectome dysfunctions were corrected in the trained group, as the averaged connectivity maps of the trained rats showed very similar patterns of co-activation with the EC between genotypes. Moreover, in trained rats, no significant differences between genotypes were observed in the statistical map, suggesting that cognitive stimulation prevents this network deterioration (Fig. 3b).

**Figure 3:**
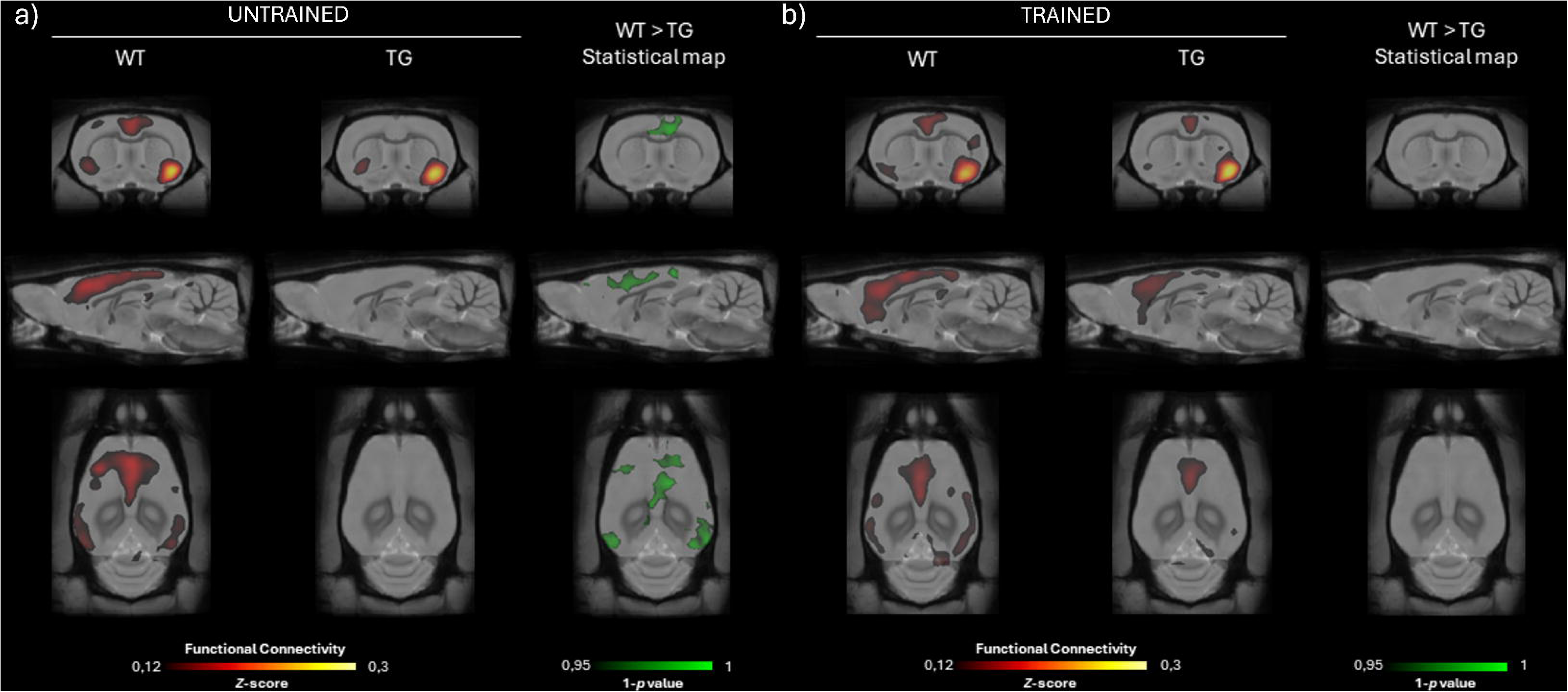
Seed-based analysis of the EC shows restored functional connectivity in trained TG rats. Seed based analyses were performed on the left EC. Average functional connectivity maps per genotype (left and centre) and statistical map indicating significant family-wise error-corrected differences (right) for a) untrained and b) trained rats (WT > TG; p<0.05, FWE-corrected). Image left is subject’s right.

### 3.3. Cognitive stimulation prevents recognition memory deficits in 19-months-old male TG rats

Since a protective effect of cognitive stimulation was observed in the functional connectivity of trained rats, the working memory of the rats was evaluated to determine if this positive effect extended to cognitive function. The NOR test was used to evaluate recognition memory as it relies on the innate bias of rodents to explore novel stimuli, indicating that a representation of the familiar object exists in the animal’s memory. The NOR test is widely used to evaluate hippocampal-dependent working memory in experimental models of AD, given that it is this type of memory that deteriorates in dementias, including AD (39). At 19-months-old, including both sexes, a two-way ANOVA revealed a significant effect of treatment (p<0.05), where trained rats had a better RI, and a close to significant effect of genotype (p=0.05) (Fig. 4a). When including the effect of sex in the analysis, a three-way ANOVA revealed a significant effect of treatment (p<0.05) and close to significant effect of genotype (p=0.05) (Fig. 4b). Training particularly improved the RI of male rats, with a significant improvement in male TG rats (p<0.05).

**Figure 4.**
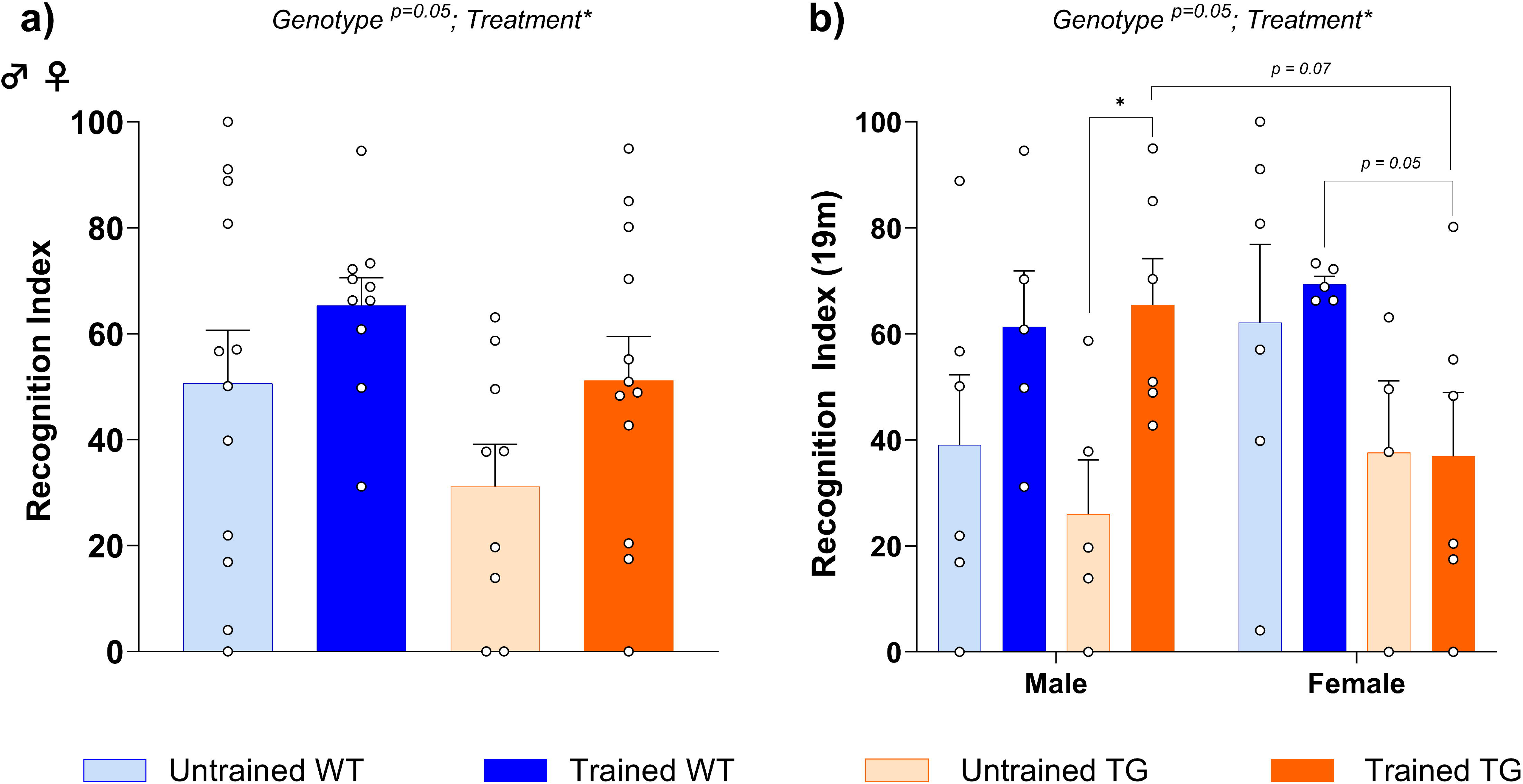
Cognitive stimulation preserves memory function in aged male TG rats. a) Recognition memory function was assessed in 19-month-old rats using the RI, the time spent in novel object relative to total object exploration. A RI greater than 50% indicates a novelty preference. The effects of genotype and treatment and the interactions between these factors were assessed by a two-way ANOVA with Tukey post-hoc tests; n= 9-12 per group when sexes combined. b) RI was also analysed separately by sex, where the effect of genotype, treatment and sex was assessed by a three-way ANOVA test with Fisher’s LSD post-hoc analyses. N= 4-6 per experimental group. Data is presented as mean ± SEM; * p<0.05; ** p<0.01; *** p<0.001.

Of note, a significant effect of training (p<0.05) was also observed in the exploration time during the NOR test, where trained rats explored more compared to untrained ones (Supp. Fig. 1).

### 3.4. Cognitive stimulation increased NeuN^+^ neuronal density in the EC of female WT rats

To evaluate whether connectivity preservation after cognitive training was due to the preservation of neurons, we counted the relative number of NeuN^+^ neurons in the primary regions implicated in recognition memory (Fig. 5a). Three-way ANOVAs revealed a significant sex x genotype x treatment interaction in the EC (p<0.01) (Fig. 5b), a significant effect of genotype in the DG (p<0.05) (Fig. 4e) and a close to significant effect of genotype in the CA1 (p=0.07) (Fig. 5c). Subsequent post-hoc analyses revealed that in the CA1, male untrained TG rats had significantly fewer NeuN^+^ neurons compared to WT littermates (p<0.05) (Fig. 5c). Moreover, female trained WT rats exhibited significantly higher numbers of NeuN^+^ neurons in the EC compared to untrained littermates (p<0.01), female trained TG rats (p<0.01) and male trained WT rats (p<0.01).

**Figure 5.**
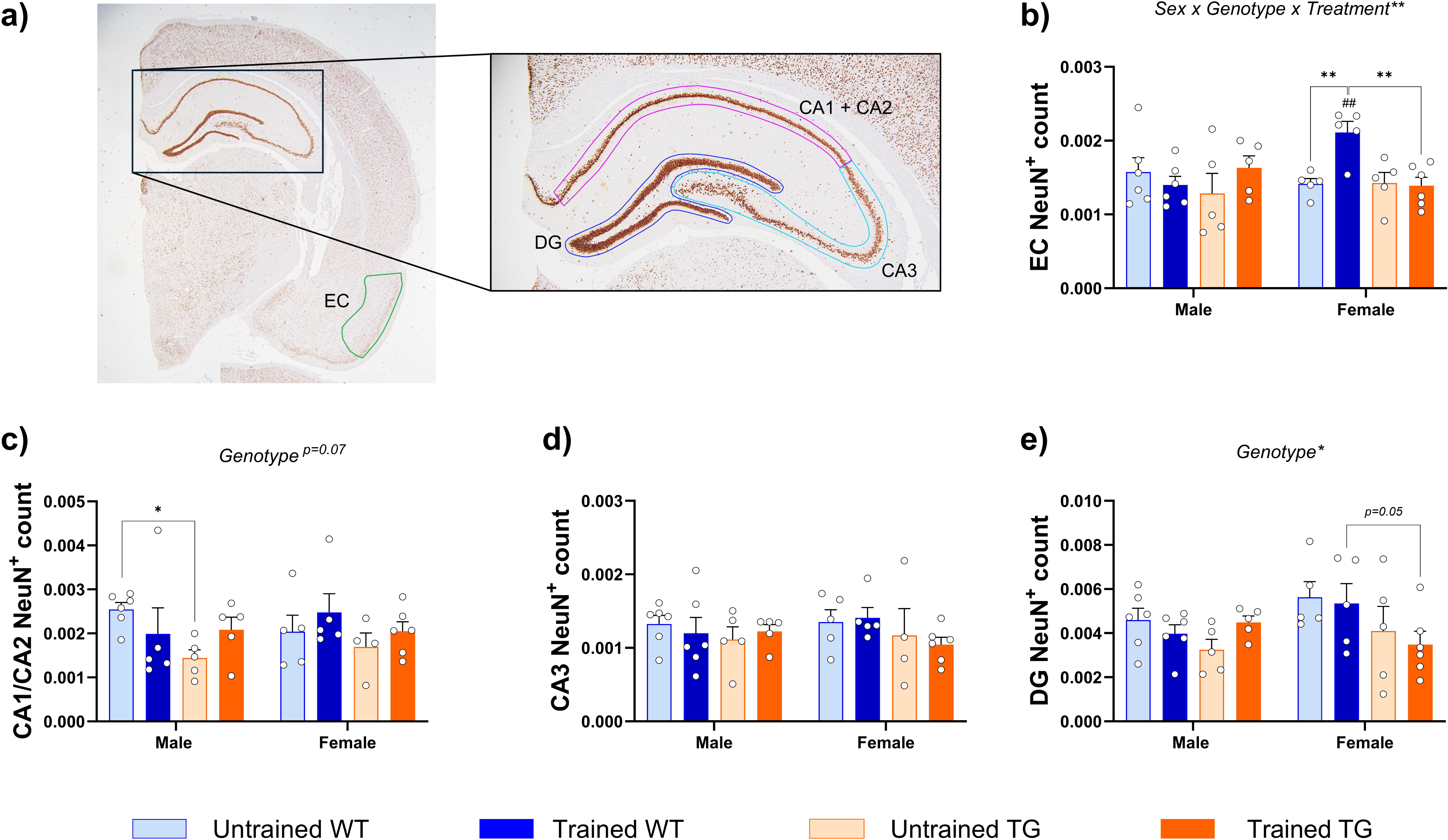
Relative NeuN^+^ count in 19-month-old rats. a) Representative immunohistochemistry image of NeuN^+^ cells (brown). For the b) EC c) CA1/CA2 d) CA3 and e) DG the number of NeuN^+^ neurons was obtained and normalised to its respective ROI to obtain a NeuN^+^ count per region. The effects of sex, genotype and treatment were assessed by a three-way ANOVA followed by Fisher’s LSD test. N= 4-6 per experimental group. Data is presented as mean ± SEM; * p<0.05; ** p<0.01; *** p<0.001.

### 3.5. Cognitive stimulation significantly decreased the number of PV^+^ interneurons in the DG

We also evaluated the PV^+^ interneuron count in the CA1, CA3 and DG of the hippocampus (Fig. 6a). Three-way ANOVAs revealed a significant effect of genotype in the CA3 (p<0.05), observing fewer PV^+^ cells in TG rats compared to WT (Fig. 6c). Moreover, a significant effect of treatment was observed in the DG (p<0.01), where trained rats (both WT and TG) showed significant less PV^+^ interneurons (Fig. 6d). Subsequent post-hoc analyses revealed a significant decrease in PV^+^ interneurons in the DG of male trained WT rats (p<0.05) compared to their untrained littermates. A similar trend was observed in the remaining groups.

**Figure 6.**
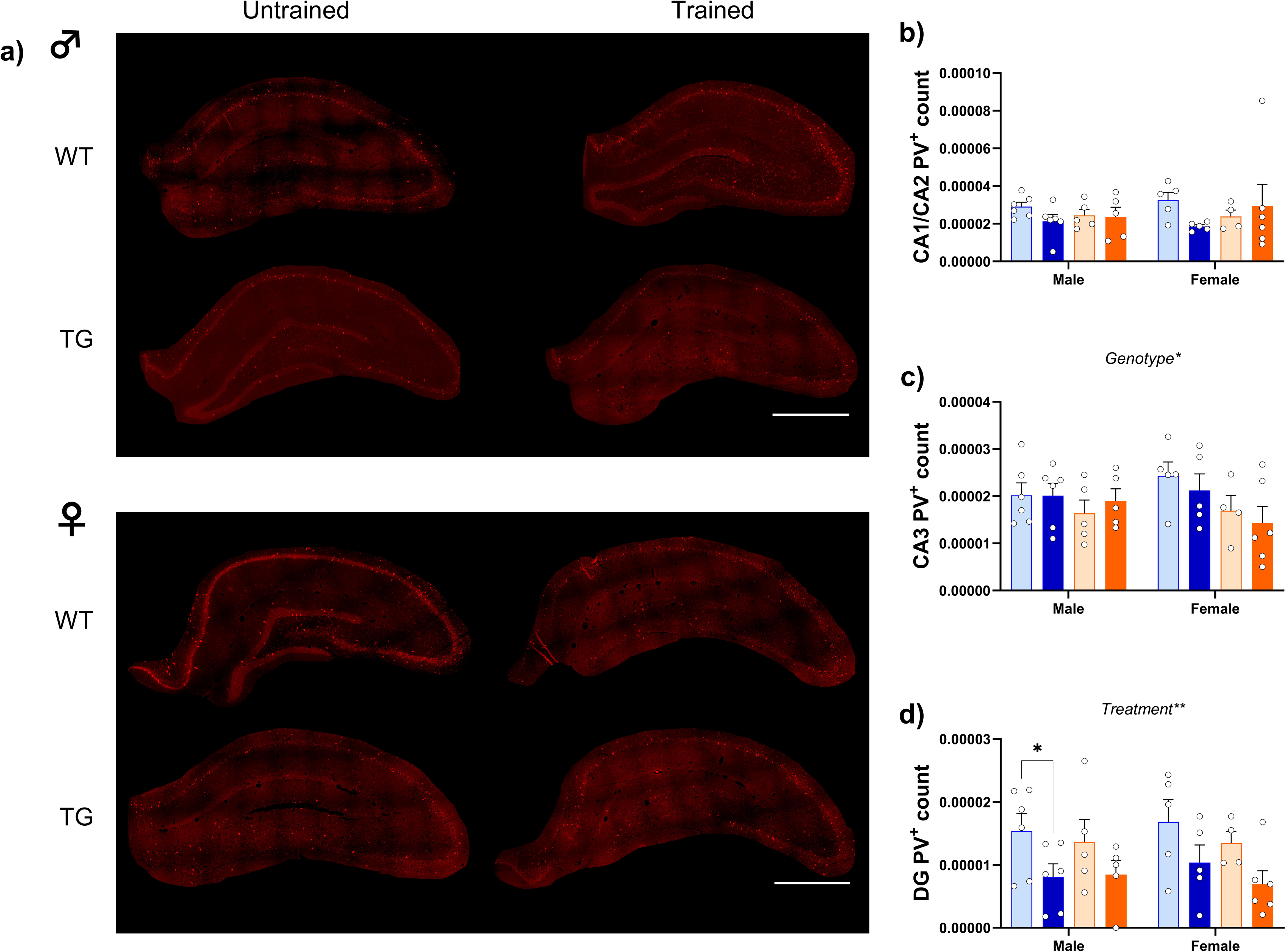
Relative PV^+^ count in 19-month-old rats. a) Representative immunofluorescence image of PV^+^ interneurons in the DG (red). For each experimental group, PV^+^ interneuron numbers in the b) CA1/CA2 c) CA3 and d) DG were obtained and normalised to the area of each region. The effects of sex, genotype and treatment were assessed by a three-way ANOVA followed by Fisher’s LSD test. Scale bar represents 1 mm. N= 4-6 per experimental group. Data is presented as mean ± SEM; * p<0.05; ** p<0.01; *** p<0.001.

### 3.6. Cognitive stimulation restored the expression of neuroplasticity proteins in male TG rats

To investigate whether cognitive stimulation was contributing to preserve neuronal plasticity, the levels of synaptic proteins, such as PSD95, TrkB, VGLUT, RPS6 and S6, were assessed in the cortex of 19-month-old rats (Fig. 7 and Supp. Fig. 2). The lysates included the motor, somatosensory, auditory, insular and entorhinal cortices. Corresponding immunoblots can be found in the supplementary material.

**Figure 7.**
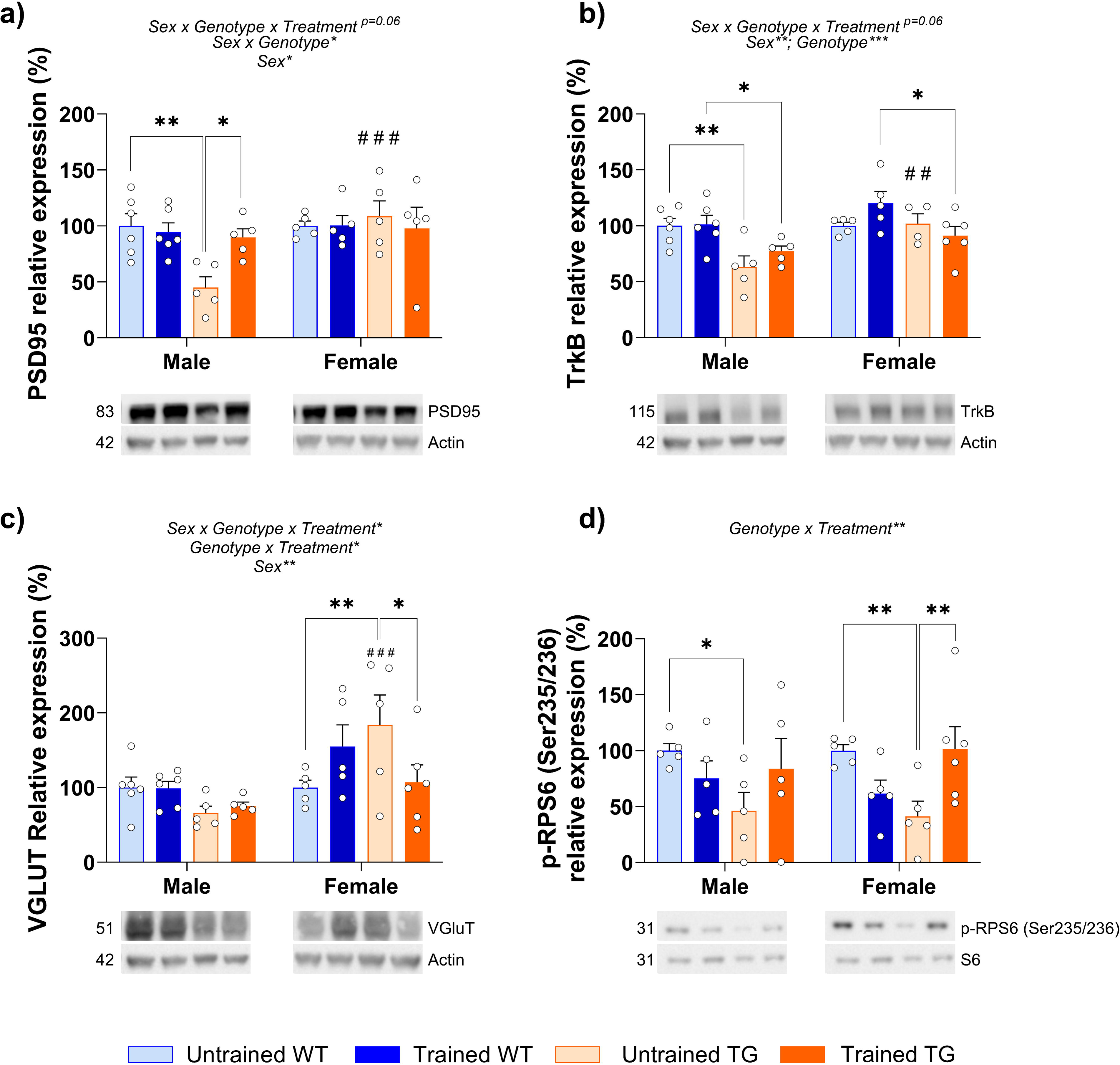
Cognitive stimulation prevents altered protein expression in TG rats. a) Immunoblottings and densitometric quantifications of PSD95 normalised to actin, expressed as a percentage relative to the untrained WT conditions. b) Immunoblottings and densitometric quantifications of TrkB normalised to Actin, expressed as a percentage relative to the untrained WT conditions. c) Immunoblottings and densitometric quantifications of VGLUT normalised to actin, expressed as a percentage relative to the untrained WT conditions. d) Immunoblottings and densitometric quantifications of p-RPS6 normalised to s6, expressed as a percentage relative to the untrained WT conditions. Respective molecular weight in kDa. The effects of genotype, treatment and sex and the interactions between these factors were assessed by a three-way ANOVA followed by Fisher’s LSD test. N= 5-6 per experimental group. Data is presented as mean ± SEM; * p<0.05; ** p<0.01; *** p<0.001; Significant differences between male and females per experimental group are denoted with “#”.

Three-way ANOVAs revealed a significant interaction between sex x genotype x treatment in the relative expression of VGLUT (p<0.05) (Fig. 7c) and a close to significance interaction in the relative expression of PSD95 and TrkB (p=0.06, both) (Fig. 7a and 7b). A significant genotype x treatment interaction in the relative expression of VGLUT (p<0.05) and p-RPS6 (p<0.01) (Fig. 7c and 7d). Moreover, a significant effect of sex was found in PSD95, TrkB and VGLUT (p<0.05, p<0.01 and p<0.01, respectively) (Fig. 7a, 7b and 7c) and a significant effect of genotype in TrkB (p<0.001) (Fig. 7b). Post-hoc analyses revealed basal sexual dimorphisms in untrained TG rats, where female rats had significantly higher expression of PSD95, TrkB and VGLUT compared to males (p<0.001, p<0.01 and p<0.001, respectively) (Fig. 7a, 7b and 7c). Male untrained TG rats had lower expression of key neuroplasticity markers, including PSD95, TrkB and p-RPS6 compared to male untrained WT rats (p<0.01, p<0.01 and p<0.05, respectively) (Fig. 7a, 7b and 7d). Cognitive training restored the protein expression of PSD95 and p-RPS6 in TG rats to a WT-like protein expression. Female untrained TG rats had lower expression of p-RPS6 compared to untrained WT littermates (p<0.01) and trained TG rats had significantly higher expression of p-RPS6 compared to untrained littermates (p<0.01) (Fig. 7d), restoring p-RPS6 back to a WT-like expression level. In summary, untrained rats displayed sexual dimorphisms in both their protein profile and response to cognitive stimulation. Female untrained TG rats had a higher expression of PSD95, TrkB and VGLUT compared to male littermates and cognitive stimulation normalised the expression of PSD95 in male TG rats and p-RPS6 in both sexes.

### 3.7. Cognitive stimulation increases the presence of microglia around A**β** plaques in 11-month-old TG rats

Microglia cells have a neuroprotective role in amyloid β-protein clearance and they cluster around Aβ plaques forming barriers to prevent plaque expansion and further glial activation (40). For this reason, we first investigated whether cognitive stimulation reduced Aβ burden in TG rats. However, at 18-months-old, we only found a close to significant effect of treatment in the DG (p=0.05) (Supp. Fig. 3). Next, we addressed the impact of cognitive training on the density of microglial cells surrounding Aβ plaques. Data were collected at two time points (at 11 and 18 months of age). The sample size at 11 months was too small to allow for sex-specific analyses, so the data is presented with sexes combined. Two-way ANOVAs were conducted in the EC, DG, CA1 and CA3 (Fig. 8a, b, c and d). A significant interaction of treatment and age was observed in the EC (p<0.05), as well as a significant effect of treatment (p<0.01) (Fig. 8a). A significant effect of age was observed across all the studied areas (p<0.001, for all) (Fig. 8a, b, c and d).

**Figure 8.**
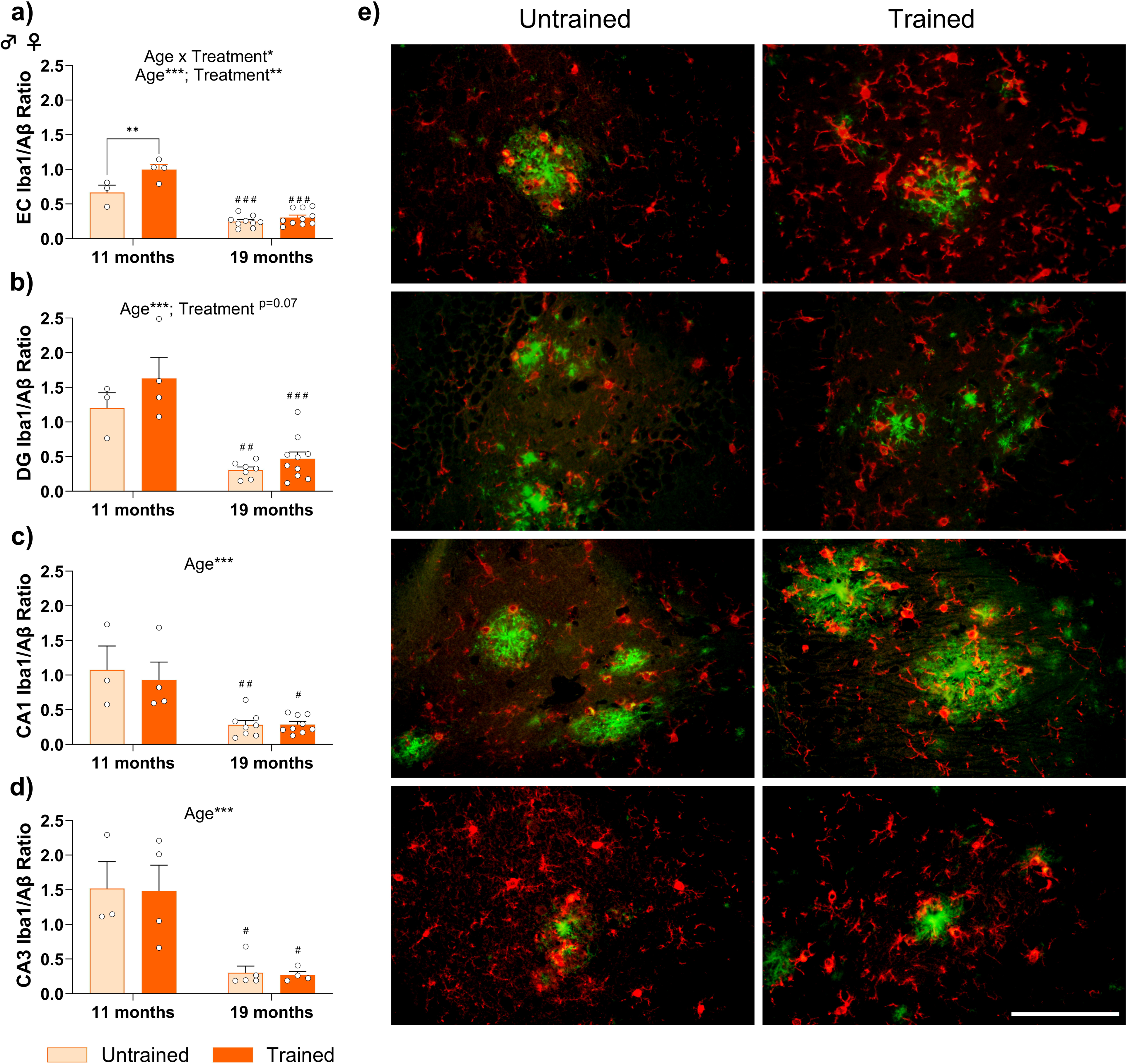
Microglia-to-plaque size ratio significantly decreases as TG rats age, independently of treatment. Mean ratio of microglia-to-plaque size in the a) EC b) DG c) CA1 and d) CA3 per rat, in 11- and 19-month-old TG rats. The effects of treatment and age and the interactions between these factors were assessed by a two-way ANOVA with Tukey post-hoc tests. Scale bar represents 50 µm. At 11-month-old n= 3-4 and at 19-months old n= 9-10 per experimental group. Data is presented as mean ± SEM; * p<0.05; ** p<0.01; *** p<0.001; Significant differences between 11 and 19 months per experimental group are denoted with “#”. b) Representative double-immunofluorescent staining of microglia (Iba1^+^ in red) and plaques (Aβ^+^ in green) for each corresponding area in 11-month-old rats.

In 11-month-old rats, a tendency was observed where trained TG rats had a greater microglial density relative to Aβ plaques, compared to untrained rats, as observed in Fig. 8b. Post-hoc Tukey test revealed the increase in microglial density was statistically significant in the EC (p<0.01) (Fig. 8a). Compared to younger rats, 19-month-old rats had a significantly lower density of microglia cells around Aβ plaques in all brain regions, irrespective of training (Untrained: EC, p<0.001; DG, p<0.01; CA1, p<0.01, CA3, p<0.05; and trained: EC, p<0.001; DG, p<0.001; CA1, p<0.05 and CA3, p<0.05) (Fig. 8a, b, c and d). Furthermore, by 19 months of age the significant effect of cognitive training was no longer observed. Thus, cognitive stimulation enhanced the proportion of microglia surrounding plaques, particularly in the EC, in an age-dependent manner as this effect of training was no longer observed in older rats. Independently of treatment, by 19 months, there was a drastic decrease in microglia per Aβ plaque, suggesting that microglial function is compromised with ageing.

### 3.8. Pre-amyloid cognitive stimulation transiently preserves a WT-like microglial morphology in 11-month-old TG rats

Since the EC is involved in successful DNMS performance (15,41) we investigated whether differences in microglia morphology (circularity and branch number) were detectable between TG rats and WT rats and whether pre-amyloid cognitive stimulation impacted microglial morphology in this region, as it is often used as a proxy for phenotype (Fig. 9a) (42).

**Figure 9.**
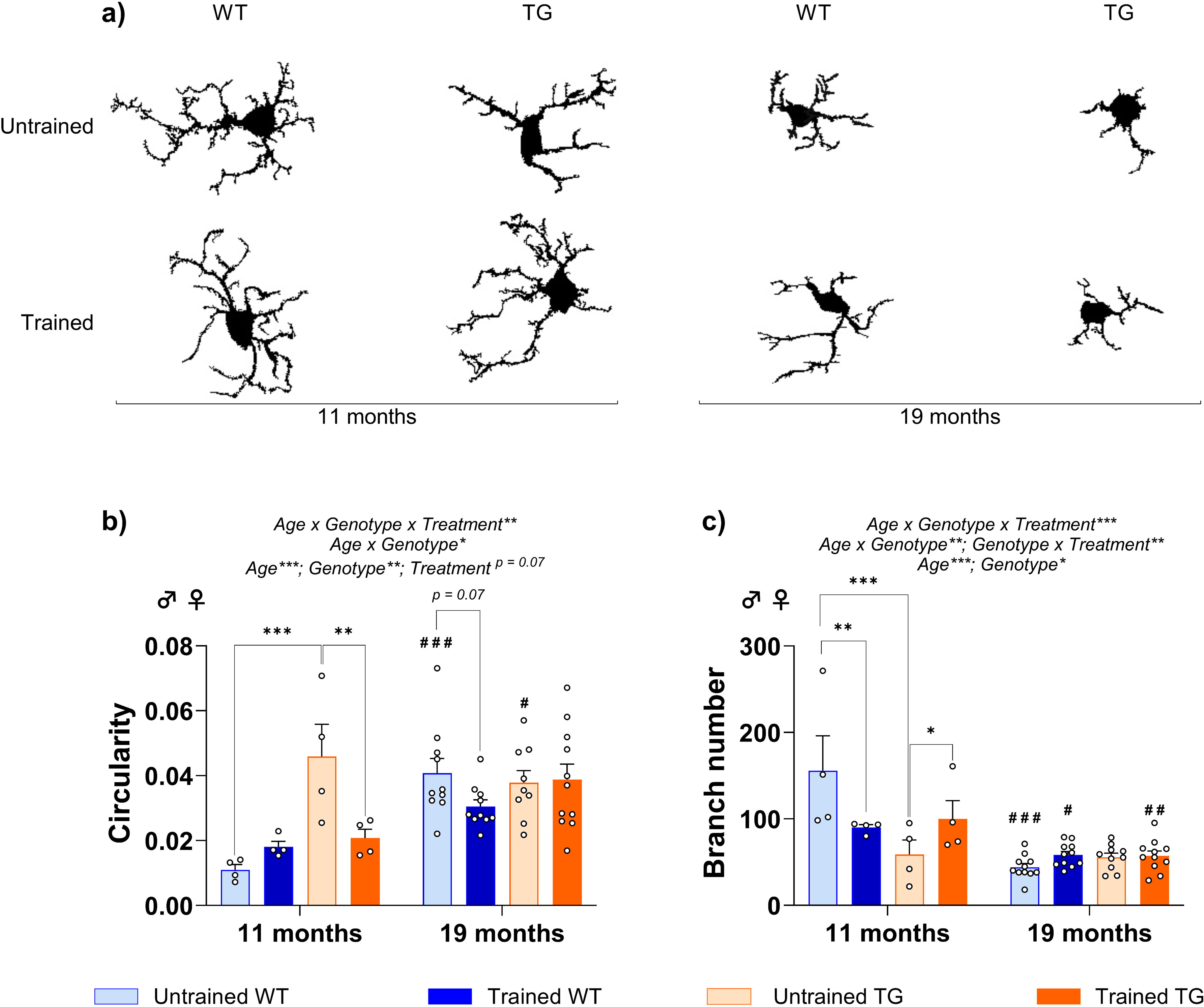
Cognitive stimulation protects microglial morphology in 11-month-old TG rats. a) Representative binary images of EC microglia from 11- and 19-month-old rats per experimental group, obtained by confocal microscopy and cleaned with ImageJ software. Mean b) circularity measured as (4π x area) ÷ perimeter^2^ and c) microglial branch number per rat. The effects of genotype, treatment and age and the interactions between these factors were assessed by a three-way ANOVA followed by Fisher’s LSD test. At 11-month-old, n= 4 and at 19-month-old n= 10-11 per experimental group. Data is presented as mean ± SEM; * p<0.05; ** p<0.01; *** p<0.001; Significant differences between 11 and 19 months per experimental group are denoted with “#”.

Including both sexes, three-way ANOVA revealed a significant age x genotype x treatment interaction (circularity, p<0.01 and branch number, p<0.001), a significant interaction between age x genotype (circularity, p<0.05 and branch number, p<0.01), a significant interaction between genotype and treatment (p<0.01, branch number) and a significant effect of age (p<0.001, both) and genotype (circularity, p<0.01 and branch number, p<0.05) (Fig. 9b and c). Post-hoc analyses revealed that in 11-month-old rats, the microglia in untrained TG rats was more circular and less branched (p<0.001, both) (Fig. 9b and c), suggesting a more reactive phenotype, compared to untrained WT littermates. Trained TG rats had significantly less circular and more branched microglia compared to untrained littermates (p<0.01 and p<0.05, respectively) (Fig. 9b and c), resembling a WT-like phenotype. As rats aged, microglia became significantly more circular (WT untrained: p<0.001 and TG untrained: p<0.05) and less branched (WT untrained: p<0.001; WT trained: p<0.05 and TG trained: p<0.01) (Fig. 9b and c). In aged 19-month-old rats, the effect of pre-amyloid cognitive stimulation was lost as no significant effect was found between treatment group. These results suggest that pre-amyloid cognitive intervention transiently preserves the microglial morphology in 11-month-old TG rats to resemble that observed in WT rats, as these were less circular and more branched. As rats aged, microglia become more round and less branched, and the effect of training dissipated.

## 4. Discussion

Our study uniquely examines the long-term impact of early cognitive stimulation on both healthy and pathological aging, while also accounting for sex differences, in an AD rat model. Promoting healthy aging involves mitigating cognitive decline, enhancing brain function, and strengthening CR (43). Following established research guidelines for studying CR (7), our longitudinal study demonstrates that pre-amyloid cognitive stimulation preserves whole-brain functional connectomics and in particular EC connectivity, leading to improved memory, particularly in male TG rats. This protective effect may be attributed to increased expression of neuronal plasticity markers and a less reactive neuroinflammatory profile in trained rats.

A well-organized modular brain structure, in particular higher segregation of functional networks, is known to support cognitive resilience and delay cognitive decline in AD (44). Additionally, higher resting-state global connectivity correlates with greater CR in patients with mild cognitive impairment (45). Despite early alterations in DWI-based structural connectomics, we previously showed that whole-brain functional connectivity in male TG animals remained unaltered until later disease stages (12,13). Within the same dataset, resting-state networks particularly the somatosensory and anterior components of the default mode network exhibited different connectivity temporal patterns in TG rats compared to WT animals (14). Other studies employing dynamic rs-fMRI analysis identified early-stage connectivity reductions in the basal forebrain and regions within the default mode-like network as early as 4 months of age, during the pre-plaque stage (46,47). However, these studies were conducted exclusively in male TG and WT subjects. In contrast, a longitudinal study focusing solely on females reported reduced functional connectivity across brain regions, with the most severe phenotype observed at 10 months of age (48). The differences in analytical approaches across these studies likely contribute to the variation in findings. However, a key limitation remains: none of these studies included both males and females.

We employed whole-brain network analyses, which have been proposed to bridge phylogenetic differences between rodents and humans in translational neuroscience (49). Considering the sex-specific prevalence of AD and documented differences in cognitive resilience between men and women (50), we included both sexes in our experimental design. Our findings demonstrate that whole-brain connectomics metrics, describing network integration (global efficiency) and network segregation (clustering coefficient) (26), are influenced by early cognitive stimulation in a sex-dependent manner. While early stimulation enhanced these metrics in males from 3 to 15 months of age, females did not exhibit the same temporal protective effect on whole-brain functional connectomics. However, females displayed a significant age-genotype interaction that was not observed in males, suggesting that network alterations in female TG animals could not be fully compensated by early cognitive stimulation. In line with this, a functional connectome study in dementia patients reported that females experience stronger dementia-related changes in key brain network regions compared to males (51). Additionally, evidence suggests that AD pathology may have a greater impact on brain structure in women, indicating lower brain resilience in females with a clinical diagnosis (50).

Given its pivotal role in memory processing and communication with the hippocampus and cortex, investigating EC connectivity is essential. The EC is one of the earliest and most severely affected regions in AD, showing atrophy, neuronal loss, and synaptic degeneration (52,53). Structural changes in AD, including reductions in brain volume, dendritic spines, and synapse densities, are particularly evident in the EC, hippocampus, and cerebral cortex (54,55). Moreover, disruptions in EC connectivity contribute to cognitive decline making it a crucial target for understanding disease progression, resilience mechanisms, and therapeutic interventions (38). In this study, we assessed EC connectivity in aged TG and WT animals at 18 months, both with and without early cognitive stimulation. TG animals exhibited EC connectivity loss in key memory-related regions, including the visual, somatosensory, motor, cingulate, and retrosplenial cortices. Importantly, cognitive stimulation drastically prevented this connectivity loss, reinforcing the EC’s role as a central hub for brain resilience. Overall, these results support the hypothesis that maintaining the integrity of the brain’s functional connectivity is crucial for keeping cognitive functions.

Both the DNMS and the NOR test rely on working memory, so improvement due to DNMS training should be reflected with enhanced NOR performance, due to shared cognitive capacities (56). While the DNMS test primarily targets hippocampal-dependent memory, its benefits also extend beyond to cortical memory processes, as evaluated by the NOR test (19). By 18 months of age, WT rats are considered old and exhibit normal aging-related working memory deficits (20,57), while TG rats show significant impairments in memory as early as 12 months (11,58–60). Accordingly, we observed that 18-months-old untrained rats could not discriminate between the novel and familiar object, regardless of genotype. Importantly, we showed that cognitive stimulation improved the RI in both male TG and WT rats. Although the overall impact of cognitive stimulation on recognition memory was limited, the significant improvement shown in TG males supports the beneficial effects of cognitive stimulation in healthy and pathological ageing. In agreement with our findings, periodic cognitive enrichment improved the memory of 18-month-old 3xTg-AD mice when compared to naïve littermates (19). Similarly, exercise training had protective effects in 18-month-old male TG rats when assessing spatial memory with the Barnes maze task, and in 15-week-old Wistar male rats with an intrahippocampal Aβ injection as assessed by the Morris Water Maze test (61,62). These findings align with observations in humans where exercise and cognitive enrichment have been associated with preserved cognition (2). We did not observe a treatment effect in female rats where the genotype effect was more pronounced compared to males. The initial characterization of this model reported no sex differences, leading many studies to overlook the influence of sex (11). However, direct sex comparisons show phenotype differences, with female TG rats outperforming their male counterparts at 9 and 11 months in an active avoidance test for spatial memory (63) and in a delayed match-to-sample test for working memory (64).

The effect of cognitive stimulation over whole-brain network metrics, EC connectivity and recognition memory could not be explained by strong differences in hippocampal number of neurons at 18-months-old, as no treatment effects were detected. Cognitive stimulation did increase NeuN^+^ cell number in the EC of female trained WT rats, compared to untrained littermates. In this line, Chaudry et al. reported that in untreated 9-month-old rats, female TG rats had higher NeuN+ cell number in the CA1 and DG, while no sex differences were observed in the hippocampus of WT rats (65). A change in PV^+^ expression in trained rats might however contribute to the observed effect of cognitive stimulation, as a significant reduction of PV^+^ interneurons was clearly observed in the DG of trained rats (WT and TG). PV^+^ GABAergic inhibitory interneurons play a key role in maintaining the excitation-inhibition balance in the hippocampus, thereby synchronising neuronal networks during memory encoding and retrieval. Age-dependent memory loss associated with hyperexcitability has been linked to the reduction in inhibitory input from the DG to the CA3 (37). Moreover, recent findings indicate that that targeted suppression of PV^+^ interneurons in the hippocampus can enhance CA3 output to CA1, increase slow gamma power and improve memory consolidation and retrieval (66). Furthermore, Donato et al. demonstrated that PV^+^ interneurons circuits exhibit bistable states, with environmental enrichment promoting a sustained, yet reversible low-PV^+^ network configuration that is associated with enhanced structural synaptic plasticity and improved NOR. Notably, they observed that the PV^+^ count remained unchanged under different experimental conditions, but rather environmental experience impacted PV^+^ expression levels. Donato et al. also showed that PV^+^ interneuron inhibition also enhanced NOR performance (67). Collectively, this data supports the idea that the observed decrease in PV^+^ count in the DG of trained rats might be a compensatory mechanism of resilience rather than a pathological loss.

Neuroplasticity encompasses a wide range of processes, including synaptic plasticity, neuroinflammation, and neuronal connectivity (43,68). The mTOR pathway is a key regulator of memory and learning, influencing synaptic plasticity, metabolism, and protein synthesis, among other essential functions. In AD, dysregulated mTOR signalling contributes to the accumulation of protein aggregates and synaptic dysfunction, both hallmarks of disease progression (69,70). Therefore, to explore a potential neurobiological mechanism behind the observed improvement in connectivity and memory we focused on the mTOR signalling pathway, specifically an upstream regulator (TrkB) and mTOR downstream proteins (p-RPS6 and PSD-95). TrkB, the BDNF receptor, plays a crucial role in promoting neuronal survival and synaptic plasticity (71,72). PSD-95 is a post-synaptic scaffolding protein that anchors glutamate-binding receptors to excitatory post-synaptic sites, modulating synaptic strength and plasticity (72,73). S6 is a ribosomal protein phosphorylated upon activation of the mTORC1 pathway involved in synaptic protein synthesis (74). VGLUT, is a vesicular glutamate transporter, which regulates glutamate availability for synaptic transmission and is essential for synaptic plasticity (75). Our findings revealed a significant reduction in these key connectivity- and plasticity-related proteins in male untrained TG rats, compared to their WT littermates. This aligns with previous observations in AD patients, where TrkB downregulation has been linked to cognitive decline (76). Notably, cognitive stimulation reversed these deficits, restoring the expression of PSD95 in male TG rats and p-RPS6 in both sexes. Cognitive stimulation could enhance and/or optimise synaptic connections, which may partly explain why male trained TG rats had improved functional connectivity and recognition memory. Moreover, we speculate that a stronger impact of cognitive stimulation on protein expression would have been more pronounced at a younger age, considering that 18-month-old rats are in advanced stages of ageing.

Microglia are critical for shaping synaptic plasticity, and therefore connectivity, contributing to CR (68). Moreover, neuroinflammation driven by chronically activated microglia, exacerbating neurodegeneration and cognitive decline in AD (77,78). As previously described, we observed that TG rats had more circular and less branched microglia, indicative of a reactive phenotype, compared to microglia in WT littermates (65,79). Pre-amyloid cognitive stimulation had a protective, anti-inflammatory effect on 11-month-old TG rats as demonstrated by an increase in microglia-to-plaque ratio and a shift towards a homeostatic phenotype, morphologically characterised by highly ramified processes. Furthermore, environmental enrichment has also been shown to induce a protective effect on microglial morphology and prevent the inflammatory response to human amyloid β-protein oligomers, observed by higher microglial density and a more ramified phenotype (80). The anti-inflammatory effect of cognitive stimulation is relevant for its protective potential, given the mechanistical link between chronic inflammation and cognitive impairment in neurodegenerative disorders (2). We observed a significant decrease in microglial density and an ameboid phenotype in 19-month-old TG rats independently of treatment. “Inflammageing”, the accumulated effect of chronic-low grade inflammation and microglial activation with ageing (81), may override the protective effect of cognitive training. As aging has been shown to exacerbate inflammation in TG rats, resulting in hyperactive microglia by 18 months. (20). Furthermore, the dysmorphic phenotype in aged rats may reflect a senescence state, where cells loose sensitivity to their environment (82).

Given the higher prevalence of dementia in women and to compensate for the historical bias of using male subjects in preclinical research, we ensured to include both sexes in our study (83). We found that sex influenced both the progression of AD pathology and the response to cognitive stimulation. Thus, female untrained TG rats expressed significantly more PSD95, TrkB and VGLUT compared to their male counterparts, while cognitive stimulation exerted a more protective effect in male TG rats, evidenced by sustained functional connectivity, improved memory and enhanced protein expression. This sexual dimorphism in response to the cognitive stimulation may stem from sex-specific dynamics of AD pathology, as TG rats show molecular and behavioural differences in disease progression. In this regard, 9-month-old female TG rats exhibit greater Aβ plaque burden, impaired glucose metabolism and greater anxiety-like and apathic behaviour, whereas 9-month-old male TG rats display increased neuronal loss and impaired learning and memory (58,64,65,84). All these findings are in line with human research suggesting a trade-off: while women generally show a consistent cognitive advantage during healthy aging, they tend to progress more rapidly to clinical AD during pathological aging, indicating lower cognitive resilience after the onset (50). This may imply that the initial resilience observed in female rats dissipates with onset of pathology, aligning with the hypothesis that sex-specific dynamics in AD pathology and resilience contribute to distinct disease trajectories.

The main limitation of this study is the relatively small number of animals per experimental group, especially in the rats sacrificed at 11 months. This constraint stems from the complexity of our study design, which involved a longitudinal investigation including behavioural testing, MRI, brain sampling at two time points and the inclusion of both sexes. The use of more animals was also limited by the COVID-19 pandemic and cost-effectiveness considerations, particularly in terms of DNMS training hours and MRI acquisitions. Despite this limitation, the consistent tendency towards better outcomes in trained rats observed at multiscale domains support our conclusions regarding the protective effect of cognitive stimulation. Furthermore, we provide additional insights into the pathology through a sex- and time-based analysis and reemphasise the importance of understanding sexual dimorphism from a preclinical level for effective translational research. In this line, a complete longitudinal characterisation of pathology in this animal model is needed to understand the effect of sex in both the progression and response to intervention in AD.

## 5. Conclusion

In summary, our findings provide further insight into the progression of AD pathology in both female and male TG rats. We demonstrate that early periodic cognitive stimulation modifies the evolution of resting-state whole-brain functional connectomics and in particular EC connectivity, acting as a compensatory mechanism against pathological aging. In males, this notably preserves recognition memory. This protective effect might be attributed to a reduction of the inhibitory function of hippocampal PV^+^ cells, enhanced expression of plasticity-related proteins and a less reactive microglial profile in the EC. Therefore, our findings provide a potential explanation for the neuroprotective effect of cognitive stimulation and underscore the importance of early intellectual enrichment and sex-specific interventions in protecting individuals susceptible to AD.

## 6. Abbreviations

Ab: Antibodies
AD: Alzheimer’s Disease
Aβ: β-amyloid
BOLD: Blood oxygen level-dependent
BSA: Bovine serum albumin
CG: Cingulate cortex
CR: Cognitive Reserve
DAB: Diaminobenzidine
DG: Dentate Guris
DNMS: Delayed non-match to sample
FoV: Field of view
FWE: Family-Wise Error
LSD: Least significant difference
MDEFT: Modified driven equilibrium fournier transform
MRI: Magnetic resonance imaging
NOR: Novel object recognition
PBS: Phosphate-buffered saline
PlCx: Prelimbic cortex
PV: Parvalbumin
RARE: Rapid acquisition with relaxation enhancement
RI: Recognition index
ROI: Region of interest
rsfMRI: Resting state functional magnetic resonance
S.c.: Subcutaneously
SEM: Standard error of the mean
TE: Echo time
TG: TgF344-AD
TR: Repetition time
WT: Wild type

## 8. Declarations

### 8.1. Ethics approval and consent to participate

Animal work was performed in accordance with the local legislation (Decret 214/1997 of July 30th by the Departament d’Agricultura, Ramaderia i Pesca de la Generalitat de Catalunya), with the approval of the Experimental Animal Ethical Committee of the University of Barcelona, and in compliance with European legislation

### 8.2. Consent for publication

Not applicable.

### 8.3. Availability of data and materials

The datasets analysed in the current study is available from the corresponding author on reasonable request.

### 8.4. Competing interest

The authors declare that they have no competing interest.

### 8.5. Funding

This work was partially funded by Instituto de Salud Carlos III (ISCIII), PI18/00893, co-funded by the European Union “A way to make Europe” and grants PID2022-1363180B-I00 (G.S.), PID2019-107738RB-I00 (F.A.), PID2020-119236RB-I00 (C.M.) and CNS2023-144340 (G.S.) funded by MICIU/AEI/10.13039/501100011033/ and by “ERDF A way of making Europe” and “European Union NextGenerationEU/PRTR. We are indebted to the Experimental MRI 7 T Unit of the Institut d’Investigacions Biòmediques August Pi I Sunyer. CIBER-BBN is an initiative financed by the Instituto de Salud Carlos III with assistance from the European Regional Development Fund.

Also, the project has been supported by María de Maeztu Unit of Excellence (CEX2021-001159), Institute of Neurosciences of the University of Barcelona, Ministry of Science, Innovation, and Universities. Grants PREP2022-000586 (J.C.P.) and PRE2022-102993 (E.A.) were financed by MICIU/AEI /10.13039/501100011033 and by ESF+. The FI-2021 grant #FI-B-00378 was financed by the Agència de Gestió d’Ajuts Universitaris i de Recerca (AGAUR) (G.C.C.).

### 8.6. Authors’ contributions

X.L.G., E.M.M., R.T., L.M.P., F.A., C.M., A.P. and G.S., contributed to the conception and design of the study and designed the experiments. X.L.G., J.C.P., G.C.C., E.A., E.L.B., C.G.G. and G.S. performed all the experiments. F.V., J.C.P., G.C.C. and G.S. performed the statistical analysis. G.S. obtained financial support. J.C.P. and G.S. wrote the manuscript. We thank Daniel Mayans, Luis Ferreiro, Carlota Domingo, Wenhui Zhou, Irene Sánchez-Domínguez, Noelia Castedo and Xenia Sofianou for their assistance throughout the project. All authors have reviewed and corrected the manuscript; they also agreed to the published version of the manuscript.

## 8.7. Acknowledgments

We thank Daniel Mayans, Luis Ferreiro, Carlota Domingo, Wenhui Zhou, Irene Sánchez-Domínguez, Noelia Castedo and Xenia Sofianou for their assistance throughout the project. All authors have reviewed and corrected the manuscript; they also agreed to the published version of the manuscript.

## 10. Additional material

The additional material includes “Additional File 1.txt,” which contains Supplementary Figures 1–3 along with their corresponding figure legends. “Additional File 2.png” contains the corresponding immunoblots. Additionally, “Supplementary Table 1.xlsx” provides the number of rats used per experimental group, and “Supplementary Table 2.xlsx” presents the results of the statistical analyses.

## Notes

### Competing Interest Statement

The authors have declared no competing interest.

## 7. References

1. Zhang XX, Tian Y, Wang ZT, Ma YH, Tan L, Yu JT. The Epidemiology of Alzheimer’s Disease Modifiable Risk Factors and Prevention. Vol. 8, Journal of Prevention of Alzheimer’s Disease. Serdi-Editions; 2021. p. 313–21.

2. Phillips C. Lifestyle Modulators of Neuroplasticity: How Physical Activity, Mental Engagement, and Diet Promote Cognitive Health during Aging. Vol. 2017, Neural Plasticity. Hindawi Limited; 2017.

3. Carlson MC, Saczynski JS, Rebok GW, Seeman T, Glass TA, Mcgill S, et al. Exploring the Effects of an “‘Everyday’” Activity Program on Executive Function and Memory in Older Adults: Experience Corps Ò [Internet]. Vol. 48. 2008. Available from: http://gerontologist.oxfordjournals.org/

4. Park DC, Lodi-Smith J, Drew L, Haber S, Hebrank A, Bischof GN, et al. The Impact of Sustained Engagement on Cognitive Function in Older Adults: The Synapse Project. Psychol Sci. 2014;25(1):103–12.

5. Nelson ME, Jester DJ, Petkus AJ, Andel R. Cognitive Reserve, Alzheimer’s Neuropathology, and Risk of Dementia: A Systematic Review and Meta-Analysis. Vol. 31, Neuropsychology Review. Springer; 2021. p. 233–50.

6. Stern Y. Cognitive reserve in ageing and Alzheimer’s disease. Vol. 11, The Lancet Neurology. 2012. p. 1006–12.

7. Stern Y, Albert M, Barnes CA, Cabeza R, Pascual-Leone A, Rapp PR. A framework for concepts of reserve and resilience in aging. Vol. 124, Neurobiology of Aging. Elsevier Inc.; 2023. p. 100–3.

8. Stern Y, Barnes CA, Grady C, Jones RN, Raz N. Brain reserve, cognitive reserve, compensation, and maintenance: operationalization, validity, and mechanisms of cognitive resilience. Neurobiol Aging. 2019 Nov 1;83:124–9.

9. Bartrés-Faz D, González-Escamilla G, Vaqué-Alcázar L, Abellaneda-Pérez K, Valls-Pedret C, Ros E, et al. Characterizing the Molecular Architecture of Cortical Regions Associated with High Educational Attainment in Older Individuals. The Journal of Neuroscience. 2019 Jun 5;39(23):4566–75.

10. Zissimopoulos J, Crimmins E, St.clair P. The value of delaying alzheimer’s disease onset. Forum Health Econ Policy. 2015 Jan 1;18(1):25–39.

11. Cohen RM, Rezai-Zadeh K, Weitz TM, Rentsendorj A, Gate D, Spivak I, et al. A transgenic alzheimer rat with plaques, tau pathology, behavioral impairment, oligomeric Aβ, and frank neuronal loss. Journal of Neuroscience. 2013 Apr 10;33(15):6245–56.

12. Muñoz-Moreno E, Tudela R, López-Gil X, Soria G. Early brain connectivity alterations and cognitive impairment in a rat model of Alzheimer’s disease. Alzheimers Res Ther. 2018 Feb 7;10(1).

13. Muñoz-Moreno E, Tudela R, López-Gil X, Soria G. Brain connectivity during Alzheimer’s disease progression and its cognitive impact in a transgenic rat model. Network Neuroscience. 2019;4(2):397–415.

14. Tudela R, Muñoz-Moreno E, Sala-Llonch R, López-Gil X, Soria G. Resting state networks in the TGF344-AD rat model of Alzheimer’s disease are altered from early stages. Front Aging Neurosci. 2019;10(JUL).

15. Paule M, Rodriguez J. Working Memory Delayed Response Tasks in Monkeys. In 2008. p. 247–65.

16. Antunes M, Biala G. The novel object recognition memory: Neurobiology, test procedure, and its modifications. Vol. 13, Cognitive Processing. 2012. p. 93–110.

17. Takeuchi H, Taki Y, Kawashima R. Effects of Working Memory Training on Cognitive Functions and Neural Systems. Vol. 21, Reviews in the Neurosciences. 2010.

18. Burnyasheva AO, Kozlova TA, Stefanova NA, Kolosova NG, Rudnitskaya EA. Cognitive training as a potential activator of hippocampal neurogenesis in the rat model of sporadic alzheimer’s disease. Int J Mol Sci. 2020 Oct 1;21(19):1–13.

19. Yeung ST, Martinez-Coria H, Ager RR, Rodriguez-Ortiz CJ, Baglietto-Vargas D, LaFerla FM. Repeated cognitive stimulation alleviates memory impairments in an Alzheimer’s disease mouse model. Brain Res Bull. 2015 Aug 1;117:10–5.

20. Chaney AM, Lopez-Picon FR, Serrière S, Wang R, Bochicchio D, Webb SD, et al. Prodromal neuroinflammatory, cholinergic and metabolite dysfunction detected by PET and MRS in the TgF344-AD transgenic rat model of AD: A collaborative multi-modal study. Theranostics. 2021;11(14):6644–67.

21. Avants B, Tustison NJ, Song G. Advanced Normalization Tools (ANTS) Release 1.00. Insight J [Internet]. 2009 Jul 29 [cited 2025 Feb 17]; Available from: http://hdl.handle.net/10380/3113

22. Barrière DA, Magalhães R, Novais A, Marques P, Selingue E, Geffroy F, et al. The SIGMA rat brain templates and atlases for multimodal MRI data analysis and visualization. Nat Commun. 2019 Dec 1;10(1).

23. Tournier JD, Smith R, Raffelt D, Tabbara R, Dhollander T, Pietsch M, et al. MRtrix3: A fast, flexible and open software framework for medical image processing and visualisation. Vol. 202, NeuroImage. Academic Press Inc.; 2019.

24. Nilearn contributors, Chamma A, Frau-Pascual A, Rothberg A, Abadie A, Abraham A, et al. nilearn [Internet]. 2025 [cited 2025 Feb 25]. Available from: https://zenodo.org/records/14697221

25. Power JD, Plitt M, Laumann TO, Martin A. Sources and implications of whole-brain fMRI signals in humans. Neuroimage. 2017 Feb 1;146:609– 25.

26. Rubinov M, Sporns O. Complex network measures of brain connectivity: Uses and interpretations. Neuroimage. 2010 Sep;52(3):1059–69.

27. Brier MR, Thomas JB, Fagan AM, Hassenstab J, Holtzman DM, Benzinger TL, et al. Functional connectivity and graph theory in preclinical Alzheimer’s disease. Neurobiol Aging. 2014 Apr;35(4):757–68.

28. Daianu M, Jahanshad N, Nir TM, Toga AW, Jack CR, Weiner MW, et al. Breakdown of brain connectivity between normal aging and Alzheimer’s disease: A structural k-Core network analysis. Brain Connect. 2013 Aug 1;3(4):407–22.

29. Fischer FU, Wolf D, Scheurich A, Fellgiebel A. Altered whole-brain white matter networks in preclinical Alzheimer’s disease. Neuroimage Clin. 2015 Jul 27;8:660–6.

30. Sanz-Arigita EJ, Schoonheim MM, Damoiseaux JS, Rombouts SARB, Maris E, Barkhof F, et al. Loss of “Small-World” Networks in Alzheimer’s Disease: Graph Analysis of fMRI Resting-State Functional Connectivity. PLoS One. 2010;5(11).

31. Smitha KA, Akhil Raja K, Arun KM, Rajesh PG, Thomas B, Kapilamoorthy TR, et al. Resting state fMRI: A review on methods in resting state connectivity analysis and resting state networks. Vol. 30, Neuroradiology Journal. SAGE Publications Inc.; 2017. p. 305–17.

32. Winkler AM, Ridgway GR, Webster MA, Smith SM, Nichols TE. Permutation inference for the general linear model. Neuroimage. 2014 May 15;92:381–97.

33. Smith SM, Nichols TE. Threshold-free cluster enhancement: Addressing problems of smoothing, threshold dependence and localisation in cluster inference. Neuroimage. 2009 Jan 1;44(1):83–98.

34. Yushkevich PA, Piven J, Hazlett HC, Smith RG, Ho S, Gee JC, et al. User-guided 3D active contour segmentation of anatomical structures: Significantly improved efficiency and reliability. Neuroimage. 2006 Jul 1;31(3):1116–28.

35. Paxinos G, Watson C. The Rat Brain in Stereotaxic Coordinates. 1988.

36. Thacker JS, Andersen D, Liang S, Zieniewicz N, Trivino-Paredes JS, Nahirney PC, et al. Unlocking the brain: A new method for Western blot protein detection from fixed brain tissue. J Neurosci Methods. 2021 Jan 15;348.

37. Hernández-Frausto M, Vivar C. Entorhinal cortex–hippocampal circuit connectivity in health and disease. Vol. 18, Frontiers in Human Neuroscience. Frontiers Media SA; 2024.

38. Igarashi KM. Entorhinal cortex dysfunction in Alzheimer’s disease. Vol. 46, Trends in Neurosciences. Elsevier Ltd; 2023. p. 124–36.

39. Bengoetxea X, Rodriguez-Perdigon M, Ramirez MJ. Object recognition test for studying cognitive impairments in animal models of Alzheimer’s disease [Internet]. Vol. 7, Frontiers in Bioscience, Scholar. 2015. Available from: www.alz.org

40. Zhao R, Hu W, Tsai J, Li W, Gan WB. Microglia limit the expansion of β-amyloid plaques in a mouse model of Alzheimer’s disease. Mol Neurodegener. 2017 Jun 12;12(1).

41. Elliott R, Dolan RJ. Differential Neural Responses during Performance of Matching and Nonmatching to Sample Tasks at Two Delay Intervals. Vol. 19, The Journal of Neuroscience. 1999.

42. Carrier M, Robert MÈ, González Ibáñez F, Desjardins M, Tremblay MÈ. Imaging the Neuroimmune Dynamics Across Space and Time. Vol. 14, Frontiers in Neuroscience. Frontiers Media S.A.; 2020.

43. Navakkode S, Kennedy BK. Neural ageing and synaptic plasticity: prioritizing brain health in healthy longevity. Vol. 16, Frontiers in Aging Neuroscience. Frontiers Media SA; 2024.

44. Ewers M, Luan Y, Frontzkowski L, Neitzel J, Rubinski A, Dichgans M, et al. Segregation of functional networks is associated with cognitive resilience in Alzheimer’s disease. Brain. 2021 Aug 17;144(7):2176–85.

45. Franzmeier N, Duering M, Weiner M, Dichgans M, Ewers M. Left frontal cortex connectivity underlies cognitive reserve in prodromal Alzheimer disease. 2017.

46. van den Berg M, Adhikari MH, Verschuuren M, Pintelon I, Vasilkovska T, Van Audekerke J, et al. Altered basal forebrain function during whole-brain network activity at pre- and early-plaque stages of Alzheimer’s disease in TgF344-AD rats. Alzheimers Res Ther. 2022 Dec 1;14(1).

47. De Waegenaere S, van den Berg M, Keliris GA, Adhikari MH, Verhoye M. Early altered directionality of resting brain network state transitions in the TgF344-AD rat model of Alzheimer’s disease. Front Hum Neurosci [Internet]. 2024 Apr 5;18. Available from: https://www.frontiersin.org/articles/10.3389/fnhum.2024.1379923/full

48. Anckaerts C, Blockx I, Summer P, Michael J, Hamaide J, Kreutzer C, et al. Early functional connectivity deficits and progressive microstructural alterations in the TgF344-AD rat model of Alzheimer’s Disease: A longitudinal MRI study. Neurobiol Dis. 2019 Apr 1;124:93–107.

49. Pagani M, Gutierrez-Barragan D, de Guzman AE, Xu T, Gozzi A. Mapping and comparing fMRI connectivity networks across species. Vol. 6, Communications Biology. Nature Research; 2023.

50. Arenaza-Urquijo EM, Boyle R, Casaletto K, Anstey KJ, Vila-Castelar C, Colverson A, et al. Sex and gender differences in cognitive resilience to aging and Alzheimer’s disease. Vol. 20, Alzheimer’s and Dementia. John Wiley and Sons Inc; 2024. p. 5695–719.

51. Wang S, Wang Y, Xu FH, Tian X, Fredericks CA, Shen L, et al. Sex-specific topological structure associated with dementia via latent space estimation. Alzheimer’s and Dementia. 2024 Dec 1;

52. Gu NN, Li H, Cao X, Li T, Jiang L, Zhang H, et al. Different Modulatory Effects of Cognitive Training and Aerobic Exercise on Resting State Functional Connectivity of Entorhinal Cortex in Community-Dwelling Older Adults. Front Aging Neurosci. 2021 May 31;13.

53. Braak H, Braak E. Acta H’ pathologica Neuropathological stageing of Alzheimer-related changes. Vol. 82, Acta Neuropathol. 1991.

54. Khan UA, Liu L, Provenzano FA, Berman DE, Profaci CP, Sloan R, et al. Molecular drivers and cortical spread of lateral entorhinal cortex dysfunction in preclinical Alzheimer’s disease. Nat Neurosci. 2014 Feb;17(2):304–11.

55. Gó Mez-Isla T, Price JL, Mckeel DW, Morris JC, Growdon JH, Hyman BT. Profound Loss of Layer II Entorhinal Cortex Neurons Occurs in Very Mild Alzheimer’s Disease. 1996.

56. Beatty EL, Jobidon ME, Bouak F, Nakashima A, Smith I, Lam Q, et al. Transfer of training from one working memory task to another: Behavioural and neural evidence. Front Syst Neurosci. 2015 Jun 2;9(June).

57. Bizon JL, LaSarge CL, Montgomery KS, McDermott AN, Setlow B, Griffith WH. Spatial reference and working memory across the lifespan of male Fischer 344 rats. Neurobiol Aging. 2009 Apr;30(4):646–55.

58. Srivastava H, Lasher AT, Nagarajan A, Sun LY. Sexual dimorphism in the peripheral metabolic homeostasis and behavior in the TgF344-AD rat model of Alzheimer’s disease. Aging Cell. 2023 Jul 1;22(7).

59. Bernaud VE, Bulen HL, Peña VL, Koebele S V., Northup-Smith SN, Manzo AA, et al. Task-dependent learning and memory deficits in the TgF344-AD rat model of Alzheimer’s disease: three key timepoints through middle-age in females. Sci Rep. 2022 Dec 1;12(1).

60. Galloway CR, Ravipati K, Singh S, Lebois EP, Cohen RM, Levey AI, et al. Hippocampal place cell dysfunction and the effects of muscarinic M 1 receptor agonism in a rat model of Alzheimer’s disease. Hippocampus. 2018 Aug 1;28(8):568–85.

61. Yang L, Wu C, Li Y, Dong Y, Wu CYC, Lee RHC, et al. Long-term exercise pre-training attenuates Alzheimer’s disease–related pathology in a transgenic rat model of Alzheimer’s disease. Geroscience. 2022 Jun 1;44(3):1457–77.

62. Zarezadehmehrizi A, Hong J, Lee J, Rajabi H, Gharakhanlu R, Naghdi N, et al. Exercise training ameliorates cognitive dysfunction in amyloid beta-injected rat model: possible mechanisms of Angiostatin/VEGF signaling. Metab Brain Dis. 2021 Dec 1;36(8):2263–71.

63. Ndukwe K, Serrano PA, Rockwell P, Xie L, Figueiredo-Pereira ME. Brain-penetrant histone deacetylase inhibitor RG2833 improves spatial memory in females of an Alzheimer’s disease rat model. Journal of Alzheimer’s Disease [Internet]. 2025 Feb 9; Available from: https://journals.sagepub.com/doi/10.1177/13872877251314777

64. Hernandez CM, McCuiston MA, Davis K, Halls Y, Dal Zotto JPC, Jackson NL, et al. In a circuit necessary for cognition and emotional affect, Alzheimer’s-like pathology associates with neuroinflammation, cognitive and motivational deficits in the young adult TgF344-AD rat. Brain Behav Immun Health. 2024 Aug;100798.

65. Chaudry O, Ndukwe K, Xie L, Figueiredo-Pereira M, Serrano P, Rockwell P. Females exhibit higher GluA2 levels and outperform males in active place avoidance despite increased amyloid plaques in TgF344-Alzheimer’s rats. Sci Rep. 2022 Dec 1;12(1).

66. Aery Jones EA, Rao A, Zilberter M, Djukic B, Bant JS, Gillespie AK, et al. Dentate gyrus and CA3 GABAergic interneurons bidirectionally modulate signatures of internal and external drive to CA1. Cell Rep. 2021 Dec 28;37(13).

67. Donato F, Rompani SB, Caroni P. Parvalbumin-expressing basket-cell network plasticity induced by experience regulates adult learning. Nature. 2013;504(7479):272–6.

68. Verkhratsky A, Zorec R. Neuroglia in cognitive reserve. Mol Psychiatry. 2024 Dec 1;

69. Freret T, Gaudreau P, Schumann-Bard P, Billard JM, Popa-Wagner A. Mechanisms underlying the neuroprotective effect of brain reserve against late life depression. Vol. 122, Journal of Neural Transmission. Springer-Verlag Wien; 2015. p. 55–61.

70. Perluigi M, Di Domenico F, Butterfield DA. mTOR signaling in aging and neurodegeneration: At the crossroad between metabolism dysfunction and impairment of autophagy. Neurobiol Dis. 2015 Jan 29;84:39–49.

71. Xia Y, Wang ZH, Liu P, Edgington-Mitchell L, Liu X, Wang XC, et al. TrkB receptor cleavage by delta-secretase abolishes its phosphorylation of APP, aggravating Alzheimer’s disease pathologies. Mol Psychiatry. 2021 Jul 1;26(7):2943–63.

72. Dejanovic B, Sheng M, Hanson JE. Targeting synapse function and loss for treatment of neurodegenerative diseases. Vol. 23, Nature Reviews Drug Discovery. Nature Research; 2024. p. 23–42.

73. Keith D, El-Husseini A. Excitation control: Balancing PSD-95 function at the synapse. Front Mol Neurosci. 2008 Mar 28;1(MAR).

74. Meyuhas O. Ribosomal Protein S6 Phosphorylation: Four Decades of Research. Int Rev Cell Mol Biol. 2015;320:41–73.

75. Moutsimilli L, Farley S, Dumas S, El Mestikawy S, Giros B, Tzavara ET. Selective cortical VGLUT1 increase as a marker for antidepressant activity. Neuropharmacology. 2005 Nov;49(6):890–900.

76. Ginsberg SD, Che S, Wuu J, Counts SE, Mufson EJ. Down regulation of trk but not p75NTR gene expression in single cholinergic basal forebrain neurons mark the progression of Alzheimer’s disease. J Neurochem. 2006 Apr;97(2):475–87.

77. Ownby RL. Neuroinflammation and cognitive aging. Vol. 12, Current Psychiatry Reports. 2010. p. 39–45.

78. Cai Z, Hussain MD, Yan LJ. Microglia, neuroinflammation, and beta-amyloid protein in Alzheimer’s disease. Vol. 124, International Journal of Neuroscience. Informa Healthcare; 2014. p. 307–21.

79. Lier J, Streit WJ, Bechmann I. Beyond activation: Characterizing microglial functional phenotypes. Vol. 10, Cells. MDPI; 2021.

80. Xu H, Gelyana E, Rajsombath M, Yang T, Li S, Selkoe D. Environmental enrichment potently prevents microglia-mediated neuroinflammation by human amyloid β-protein oligomers. Journal of Neuroscience. 2016 Aug 31;36(35):9041–56.

81. Hanslik KL, Ulland TK. The Role of Microglia and the Nlrp3 Inflammasome in Alzheimer’s Disease. Vol. 11, Frontiers in Neurology. Frontiers Media S.A.; 2020.

82. Harry GJ. Microglia during development and aging. Vol. 139, Pharmacology and Therapeutics. Elsevier Inc.; 2013. p. 313–26.

83. Nichols E, Szoeke CEI, Vollset SE, Abbasi N, Abd-Allah F, Abdela J, et al. Global, regional, and national burden of Alzheimer’s disease and other dementias, 1990–2016: a systematic analysis for the Global Burden of Disease Study 2016. Lancet Neurol. 2019 Jan 1;18(1):88–106.

84. Lopez DC, White ZJ, Hall SE. Anxiety in Alzheimer’s disease rats is independent of memory and impacted by genotype, age, sex, and exercise. Alzheimer’s & Dementia [Internet]. 2024 Apr 16; Available from: https://alz-journals.onlinelibrary.wiley.com/doi/10.1002/alz.13813

