## Additional Files for "Pre-amyloid cognitive intervention preserves brain function in aged TgF344-AD rats, maintaining connectivity and enhancing plasticity in a sex-specific manner": Additional file 1.docx

**Supplementary material**

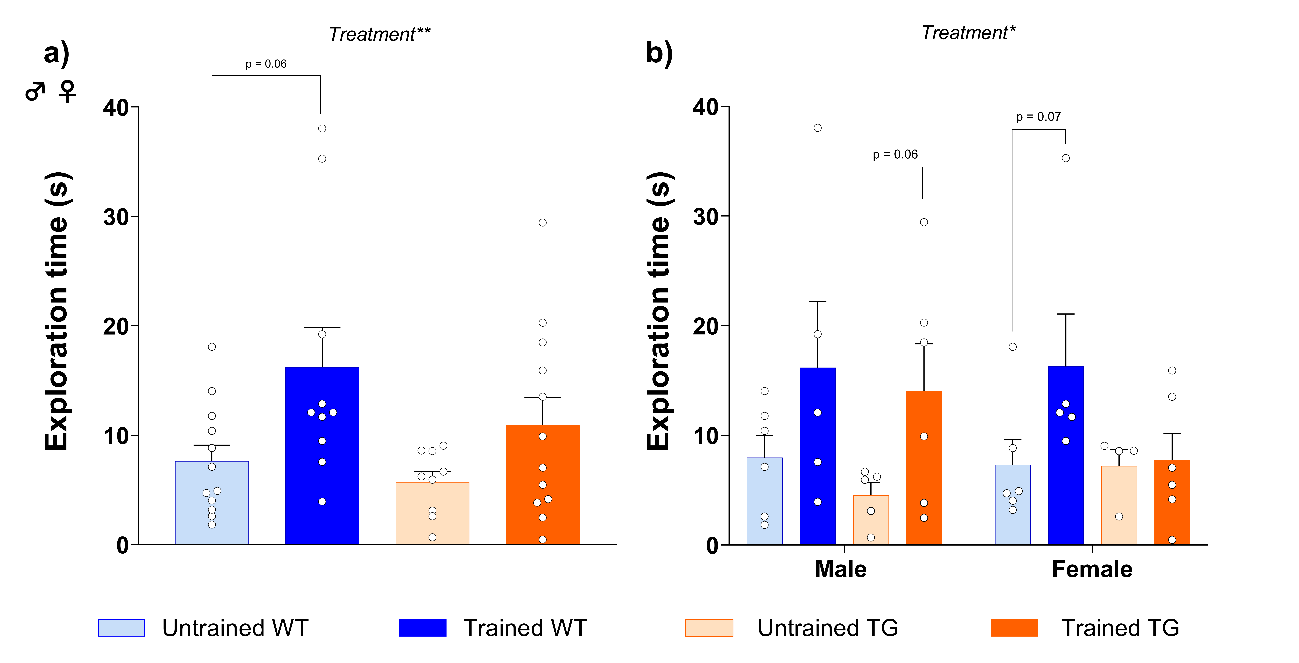

**Supplementary figure 1: Trained rats explored more during the NOR test.** A) The effects of genotype and treatment and the interactions between these factors were assessed by a two-way ANOVA test with Tukey post-hoc tests; n = 9-12 when sexes combined. b) When analysed separately by sex, where genotype, treatment and sex were assessed by a three-way ANOVA test with Fisher’s LSD post-hoc analysis. N = 4-6 per experimental group. Data is presented as mean ± SEM; * p<0.05; ** p<0.01; *** p<0.001.

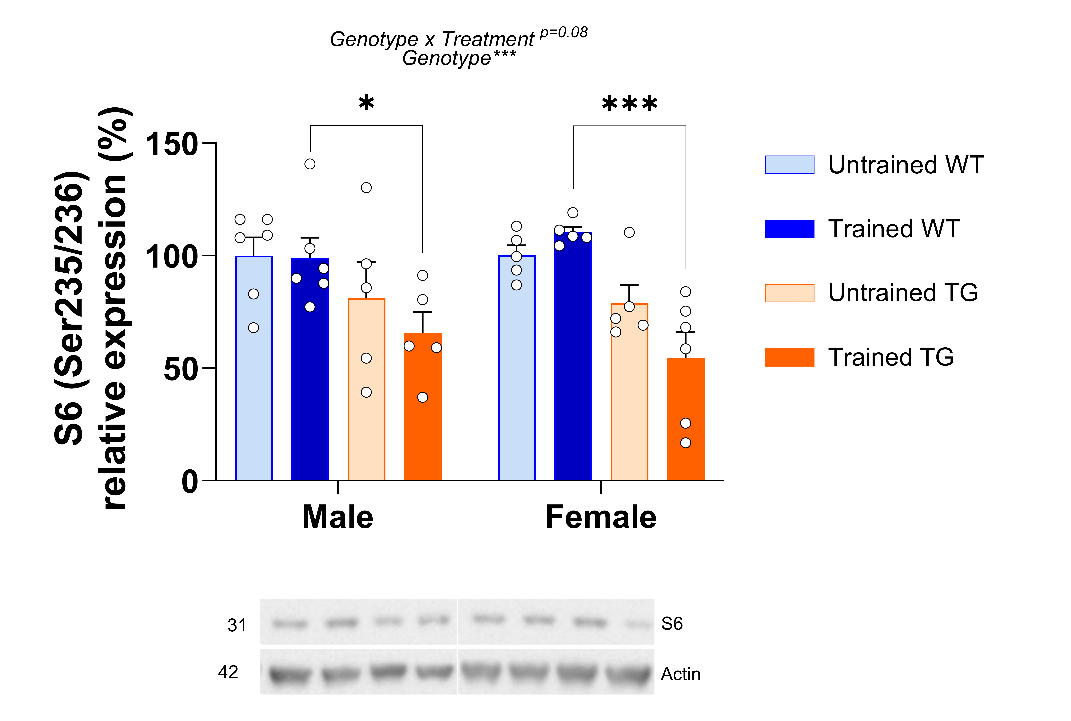

**Supplementary figure 2: Immunoblottings and densitometric quantifications of total s6** a) Immunoblottings and densitometric quantifications of S6 normalised to actin, expressed as a percentage relative to the untrained WT conditions. Respective molecular weight in kDa. The effects of genotype, treatment and sex and the interactions between these factors were assessed by three-way ANOVA test followed by Fisher’s LSD test. Data is presented as mean ± SEM; * p<0.05; ** p<0.01; *** p<0.001; Significant differences between male and females per experimental group are denoted with “#”; n= 5-6 per experimental group.

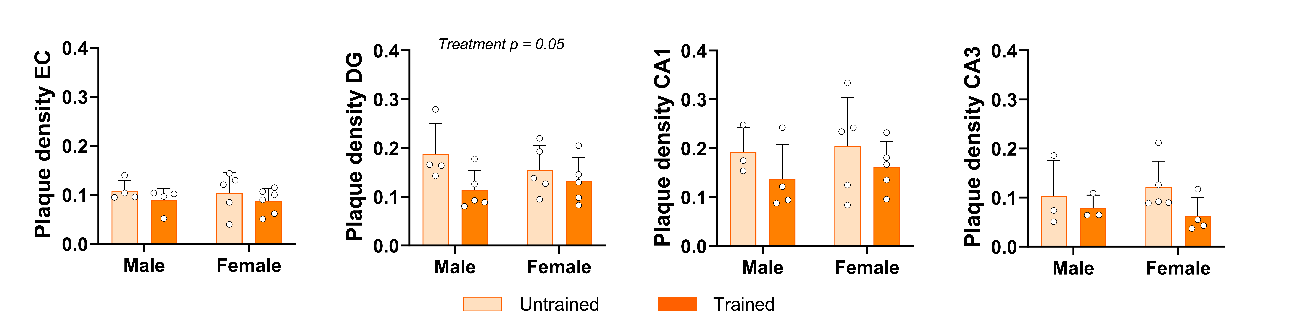

**Supplementary figure 3: Density of Aβ plaques.** The effects of treatment and sex and the interactions between these factors were assessed by two-way ANOVA with Tukey post-hoc test. Data is presented as mean ± SEM; * p<0.05; ** p<0.01; *** p<0.001; n= 6-3 per experimental group.
