## Supplementary figures and images for "Pre-amyloid cognitive intervention preserves brain function in aged TgF344-AD rats, maintaining connectivity and enhancing plasticity in a sex-specific manner"

### Additional file 2.png

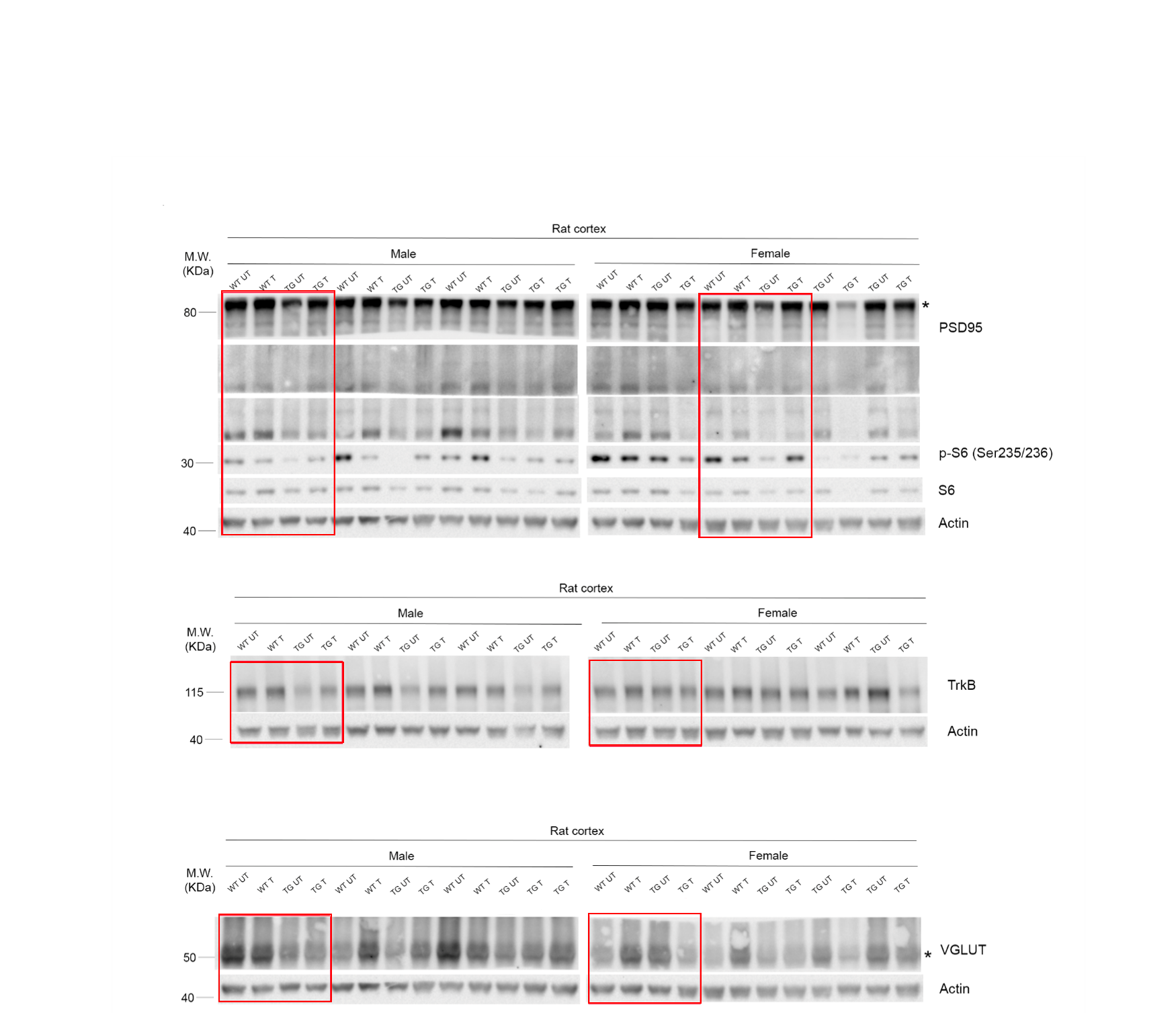
